# Exercise-induced enhancement of synaptic function triggered by the inverse BAR protein, Mtss1L

**DOI:** 10.1101/545582

**Authors:** Christina Chatzi, Gina Zhang, William Hendricks, Yang Chen, Eric Schnell, Richard H. Goodman, Gary L. Westbrook

## Abstract

Exercise is a potent enhancer of learning and memory, yet we know little of the underlying mechanisms that likely include alterations in synaptic efficacy in the hippocampus. To address this issue, we exposed mice to a single episode of voluntary exercise, and permanently marked mature hippocampal dentate granule cells that were specifically activated during exercise using conditional Fos-TRAP mice. Only a few dentate granule cells were active at baseline, but two hours of voluntary exercise markedly increased the number of activated neurons. Activated neurons (Fos-TRAPed) showed an input-selective increase in dendritic spines and excitatory postsynaptic currents at 3 days post-exercise, indicative of exercise-induced structural plasticity. Laser-capture microdissection and RNASeq of activated neurons revealed that the most highly induced transcript was Mtss1L, a little-studied gene in the adult brain. Overexpression of Mtss1L in neurons increased spine density, leading us to hypothesize that its I-BAR domain initiated membrane curvature and dendritic spine formation. shRNA-mediated Mtss1L knockdown *in vivo* prevented the exercise-induced increases in spines and excitatory postsynaptic currents. Our results link short-term effects of exercise to activity-dependent expression of Mtss1L, which we propose as a novel effector of activity-dependent rearrangement of synapses.

**One Sentence Summary:** Single episodes of voluntary exercise induced a functional increase in hippocampal synapses mediated by activity-dependent expression of the BAR protein Mtss1L, acting as a novel early effector of synapse formation.

## Introduction

The beneficial cognitive effects of physical exercise cross the lifespan as well as disease boundaries *(1, 2)*. Exercise alters neural activity in local hippocampal circuits, presumably by enhancing learning and memory through short and long-term changes in synaptic plasticity *(3, 4)*. The dentate gyrus is uniquely important in learning and memory, acting as an input stage for encoding contextual and spatial information from multiple brain regions. This circuit is well suited to its biological function because of its sparse coding design, with only a few dentate granule cells active at any one time *(5–7)*. These properties also provide an ideal network to investigate how exercise-induced changes in activity-dependent gene expression affect hippocampal structural and synaptic plasticity *in vivo.*

Most studies have focused on the cognitive effects of sustained exercise *(8–10).* However, memory improvements are well documented after acute bouts of exercise *(11, 12).* Thus, the response to a single episode of exercise may be better suited to uncover cellular and molecular cascades that form and rearrange synapses, which evolve over minutes to weeks following exercise. We reasoned that these mechanisms could be unmasked by a single period of voluntary exercise *in vivo,* followed by functional and molecular analysis of subsequent changes specific to neurons that were activated during the exercise.

## Results

cFos^cre/ERT2/Luo^ transgenic mice (Fos-TRAP) provide valid proxies of neural activity (11 and fig. S1A) and a means to permanently label activated dentate granule cells. During a two-hour exposure to running wheels, mice ran approximately 3 km. We examined activated cells 3 days post-running in Fos-TRAP mice crossed with a TdT reporter line (Fig. 1A). We used Fos immunohistochemistry at 1 hour post-exercise, confirming that this stimulates robust neuronal activity in mature granule cells (Fig. 1B). The increase in Fos expression, assessed by immunohistochemistry, matched the increase in TdTomato-positive cells (TdT^+^) measured 3 or 7 day later in Fos-TRAP mice, indicating that activated granule cells were accurately and permanently labeled during the 2 hour time window. We refer to these cells as “exercise-TRAPed”. To investigate whether a single bout of exercise activated a specific subset of granule cells, we exercise TRAPed dentate granule cells (TdT^+^) and compared this population to an ensemble activated by a subsequent re-exposure to exercise either 1 or 4 days later, as measured by Fos immunohistochemistry at 2 hours after the 2^nd^ exercise period (Fig. 1D). When the two exercise periods were separated by 24 hours, only 13% of exercise TRAPed cells were activated in the 2^nd^ exercise period. There was almost no overlap between the two neuronal ensembles (1%) when the two periods were separated by 4 days (Fig. 1D, right panel). These results indicate that our exercise protocol activated stochastic, nonoverlapping sets of granule cells, consistent with the sparse coding design of this circuit.

**Fig. 1:**
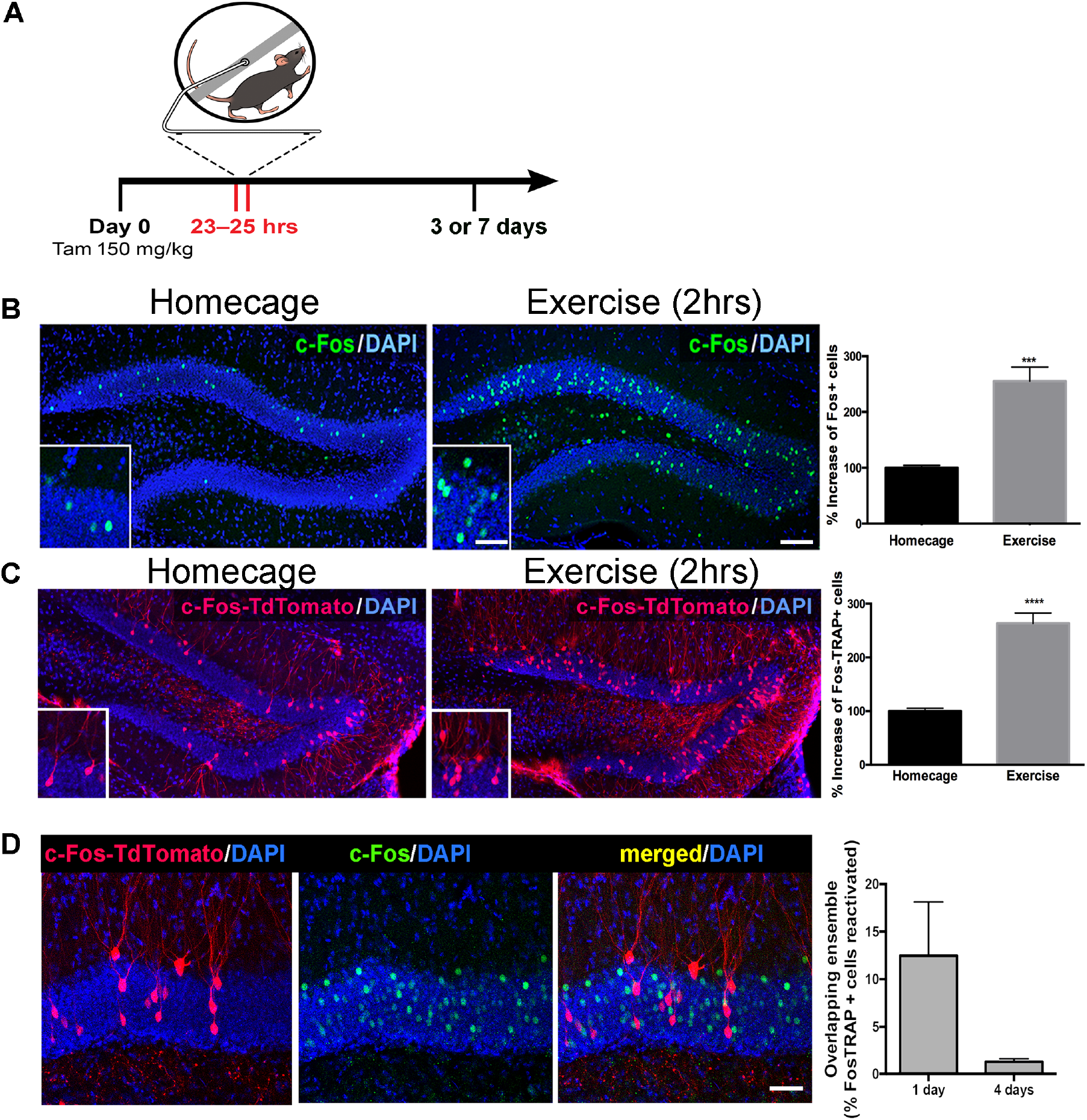
Single exposure to running wheel induces transient synaptic plasticity in exercise-TRAPed dentate granule cells. **(A)** Schematic showing exercise paradigm. Fos-TRAP:TdTomato mice were injected with tamoxifen (150 mg/kg) 24hrs before exposure to 2 hrs of voluntary exercise, while littermate controls remained in their homecage. Mice were sacrificed 3 or 7 days after exposure to the running wheel. **(B)** Voluntary exercise (2 hrs) increased neuronal activity in the dentate gyrus. Representative images of endogenous c-Fos expression in the dentate gyrus of WT mice housed in their homecage (left) or 2 hrs after exposure to voluntary exercise (middle). A single bout of exposure to exercise increased c-Fos+ cells in the dentate gyrus (right). (Homecage: 100±4 n=6, Exercise: 255±25, n=4, unpaired t-test p=0.001). Scale bars: insert 50μm, right 100μm. **(C)** Representative images of the dentate gyrus from Fos-TRAP:TdTomato mice housed in their homecage (left) or 3 days after 2hrs of voluntary exercise (middle). Voluntary exercise increased exercise-TRAPed dentate granule cells (% increase from baseline in exercise-TRAPed cells, homecage 100±5 n=5, Exercise 264±19, n=5, unpaired t-test, p<0.0001). **(D)** A single exposure to exercise tags distinct populations of activated DG granule cells due to sparse coding. (A) We used the Fos-TRAP: Tdtomato mice to tag a neuronal ensemble activated by a single exposure to exercise (2 hrs) (Tdtomato^+^). We compared these exercise-TRAPed cells to granule cells activated by a second exposure to exercise and tagged at 2 hours post-exercise using c-Fos immunohistochemistry 1 or 4 days later. Fos-TRAP:Tdtomato mice were injected with Tamoxifen (150 mg/kg) 24hrs prior to exercise. Animals were exposed to a second bout of exercise either 1 or 4 days later. (B) When the two exercise periods were separated by 24 hrs, 12.5±5.6%, (n=3) of the exercise-TRAPed cells were re-activated, whereas with a 4 day separation only 1.3±0.3% (n=4, unpaired t-test, p=0.07) overlapped, consistent with sparse labeling in the dentate gyrus.

Higher cortical information converges on the dentate gyrus through laminated perforant path axons from the entorhinal cortex. To examine whether a single exposure to exercise altered structural plasticity in exercise-activated neurons, we analyzed exercise-TRAPed cells in mice 3 or 7 days post-exercise as compared to home cage controls (Fig. 2A, left panels). Exercise-TRAPed cells showed a nearly 50% increase in dendritic spines at 3 days. This increase was limited to the outer molecular layer (OML), which interestingly, receives contextual and time information via the lateral perforant path *(13, 14),* whereas there was no change in dendritic spines in the middle molecular layer (MML), which receives spatial information via the medial perforant path *(15, 16).* Consistent with a transient response to a single stimulus, spine density in the OML returned to baseline levels by 7 days (Fig. 2A, right panels). Exercise did not affect total dendritic length in OML or MML indicating that the increase in spine density reflected an increase in the number of synapses (fig. S2).

The increase in spine density in the OML was associated with a corresponding increase in functional synaptic input in exercise-TRAPed cells (Fig. 2B). Three days after exercise, we used acute brain slices to make simultaneous whole-cell voltage clamp recordings from an exercise-TRAPed granule cell (yellow, Fig. 2B, middle) and a neighboring control granule cell (green, Fig 2B, middle). Selective stimulation of lateral perforant path (LPP) axons evoked nearly 3-fold larger EPSCs in the exercise-TRAPed cell of the simultaneously recorded cell pair (Fig. 2C,D), whereas EPSC amplitudes evoked by stimulation of medial perforant path axons (MPP) were unaffected (Fig. 2C,D). Plotting each cell pair as a function of the site of stimulation indicated that MPP cell pairs were on the unity line whereas LPP cell pairs were below the unity line, indicative of the increased EPSCs in exercise-TRAPed cells (Fig. 2E). The increased EPSC amplitudes could not be attributed to differences in intrinsic membrane properties or presynaptic release probability (fig. S3 and Table S1). These results indicate that dentate granule cells activated by a single bout of exercise show a laminar-specific increase in dendritic spines and in excitatory postsynaptic currents.

Activity-dependent gene expression is one of the main drivers of short and long-term changes in synaptic function *(17)*. To identify exercise-activated genes underlying the observed structural plasticity in the dentate gyrus, we isolated RNA from RFP^+^ nuclei of exercise-TRAPed cells (Fig. 3A,B) using laser microdissection. An ensemble of 150 individual exercise-TRAPed or adjacent non-activated (RFP^−^) mature granule cells was excised per mouse at 3 and 7 days post-exercise. Analysis of transcripts by RNASeq (fig. S4) revealed up- and down-regulated transcripts (Fig 3C, left panel, Table S2, S3) of which approximately 150 transcripts were significantly upregulated in exercise-TRAPed neurons 3 days after exercise. Gene Ontology analysis indicated that most of the upregulated transcripts had signaling and synaptic functions. The top ten upregulated genes (Fig. 3C) included several known genes involved in hippocampal function, such as the neuron-specific RNA binding protein Elavl4 and the immediate early gene Egr1 *(18, 19).* Consistent with the transient increase in dendritic spines, no transcript identified at 3 days post-exercise remained elevated at 7 days (fig. S5).

**Figure 2:**
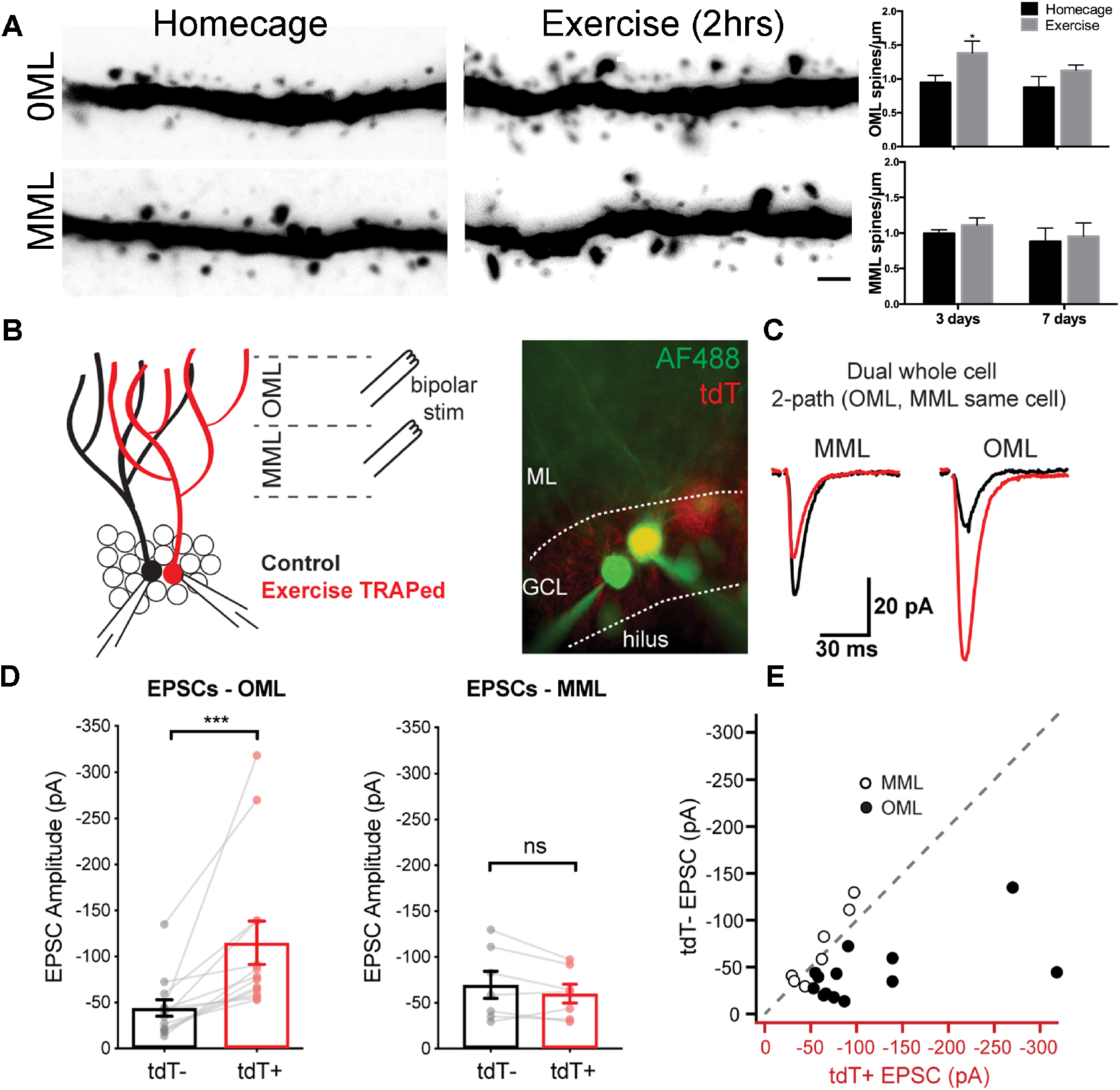
A single bout of exercise induces a laminar-specific increase in excitatory synaptic innervation. **(A)** Representative images of TRAPed granule cell dendrites in the OML and MML from mice at baseline (i.e. homecage) or 3 days post-exercise. Scale bar: 5μm. Spine densities were significantly increased in the OML of activated cells at 3 days post exercise (2-way ANOVA, OML 3 days homecage: 0.94±0.1, exercise: 1.39±0.2, p=0.02, n=5) whereas there was no difference in the OML at 7 days post-exercise (OML 7 days homecage: 0.87±0.2, exercise: 1.10±0.1, p=0.18, n=5). Spine density in the MML was unaffected (2-way ANOVA, no interaction between homecage and exercise groups, MML 3 days homecage: 0.99±0.1, exercise: 1.10±0.1; MML 7 days homecage: 0.88±0.2, exercise 0.95±0.2, n=5, p= 0.79). **(B)** Configuration for simultaneous whole-cell voltage clamp recordings of control and exercise-TRAPed dentate granule cells (left). Bipolar stimulating electrodes were placed in the OML and/or MML to activate lateral and medial perforant path axons, respectively. Cells were filled with AlexaFluor 488 dye (right) to confirm that the dendritic arbors overlapped. **(C)** Representative EPSCs from the cell pair shown at left in response to alternating MML and OML stimulation in the control (black traces) and exercise-TRAPed cell (red traces). **(D)** OML stimulation produced EPSCs that were significantly greater in exercise-TRAPed cells (EPSCs – OML amplitude: TdT^−^, −44.1 ± 8.9 pA; TdT^+^, −114.9 ± 23.5 pA; n = 13 cell pairs (8 mice); p = 0.0002, Wilcoxon matched-pairs signed rank test), whereas MML stimulation did not differ from controls (EPSCs – MML amplitude: TdT^−^ −69.7 ± 14.9 pA; TdT^+^, −60.1 ± 10.3 pA; n = 7 cell pairs (4 mice); p = 0.15, paired t-test). There was no difference in EPSC amplitudes during OML stimulation while recording from two control (both TdT^−^) granule cells (EPSC amplitudes: TdT^−^, −55.8 ± 16.3 pA; TdT^−^, −57.5 ± 13.8 pA, n = 5 cell pairs, p = 0.80, paired t-test). **(E)** EPSC amplitudes from control (y-axis) and exercise TRAPed (x-axis) from each cell pair were plotted with each point representing a cell pair. MML stimulated cell pairs (open circles) were present along the unity line (dashed line), whereas OML stimulated pairs (black circles) were shifted below unity, indicating that the larger EPSC amplitudes for exercise-TRAPed cells was specific to the OML.

**Figure 3:**
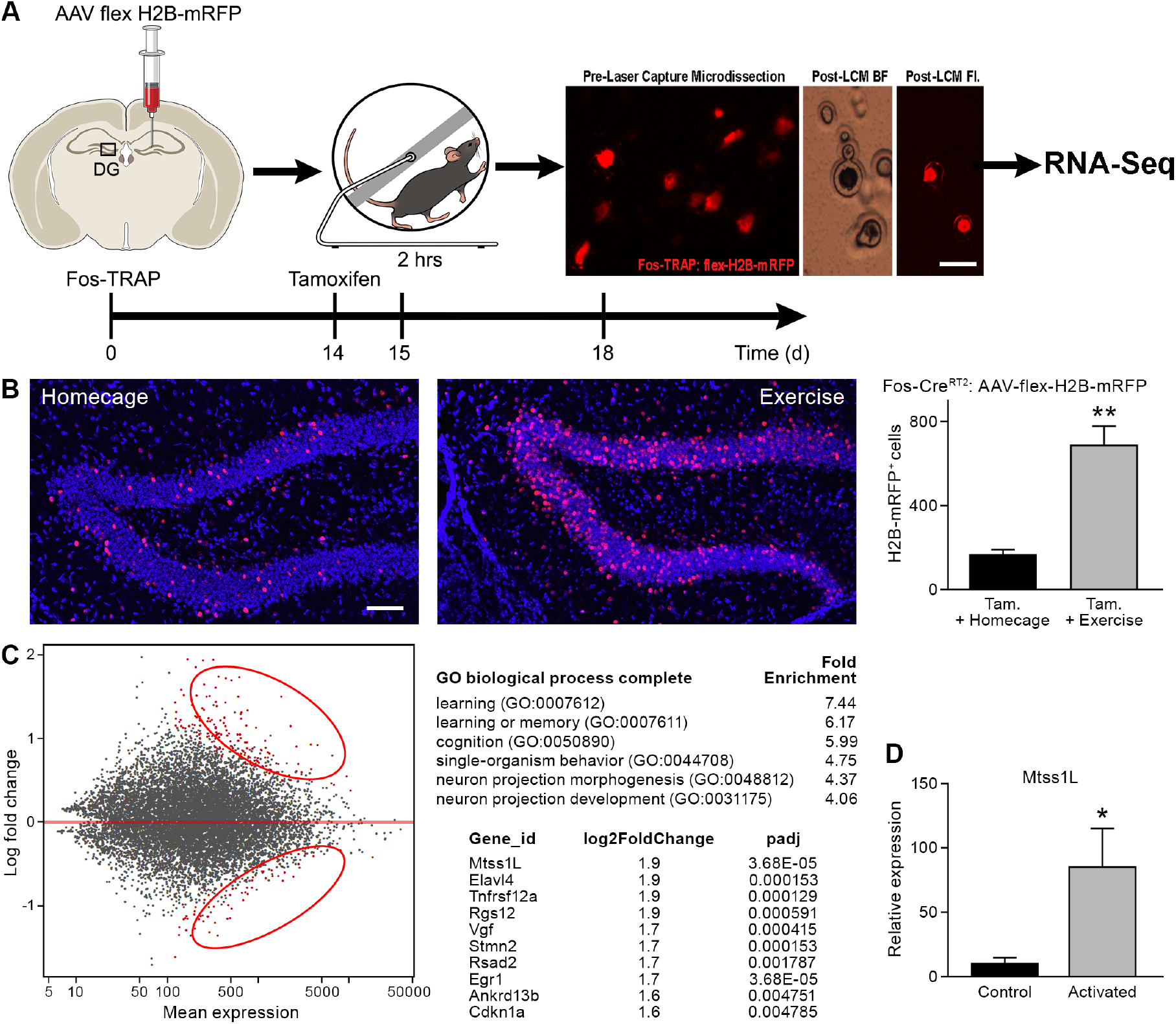
Transcriptome analysis of laser captured dentate granule cells activated by a single bout of voluntary exercise. **(A)** A virus expressing a CAG promoter followed by a loxP-flanked (‘floxed’) stop cassette-controlled Histone2B Monomeric Red Fluorescent Protein (flex H2B-mRFP), was injected stereotaxically into the dentate gyrus of Fos-TRAP mice. The nuclear tag allowed preservation of the fluorophore during laser capture microdissection. Two weeks post-viral injections, mice were injected with tamoxifen, followed 24 hours later by 2 hours of voluntary exercise. Mice were sacrificed 3 and 7 days following exercise. mRFP^+^ (exercise TRAPed) and non-activated granule cells were subsequently excised from unfixed intact tissue cryosections using laser capture microdissection, and pooled in batches of 100-150 cells per mouse. cDNA libraries were prepared and samples were processed for RNASeq library construction. Scale bar: 20μm. **(B)** Voluntary exercise increased activated mRFP^+^ cells compared to littermate controls in homecage. Scale bars: 100μm. Fos TRAPed:H2B-mRFP+ cells/50μm section was 164±26 for homecage (n=3) and 686±92 for exercise (n=4, unpaired t-test, p=0.007). **(C)** Differential expression of genes between RFP^+^ and RFP^−^ cell ensembles from 4 mice are displayed in a MA-plot (M value vs A value plot, which are Log2fold vs normalized mean expression in DEseq2), with and significantly changed (FDR <0.1, fold enrichment >2.) Upregulated and downregulated transcripts are shown as red dots (upper and lower circles, respectively). For analysis at 3 days post-exercise in exercise-TRAPed cells, the top enriched ontological clusters are listed as well as the top 10 upregulated genes. **(D)** Real-time qPCR confirmation of the Mtss1L expression in in exercise-TRAPed cells at 3 days (see also fig. S5).

We focused on Mtss1-like (metastasis-suppressor 1-like, Mtss1L), a protein that has been little studied in the adult nervous system *(20)*. Mtss1L, the most enriched transcript in our experiments, was 9-fold elevated compared to non-activated neighboring cells, as validated in exercise TRAPed cells by RT-PCR (Fig. 3D, fig. S5). To determine the spatiotemporal pattern of Mtss1L expression, we derived KOMP Mtss1L reporter mice *(21)* in which the endogenous Mtss1L promoter drives bacterial beta-galactosidase *(lacZ)* gene expression in a Cre-dependent manner. LacZ expression was undetectable in the dentate gyrus of KOMP Mtss1L^+/−^ housed in their homecage, suggesting no expression of Mtss1L under baseline conditions. In contrast, exercise-induced LacZ expression peaked at 3 days post-exercise (Fig. 4), confirming the activity-dependence of Mtss1L expression in the dentate gyrus.

**Figure 4:**
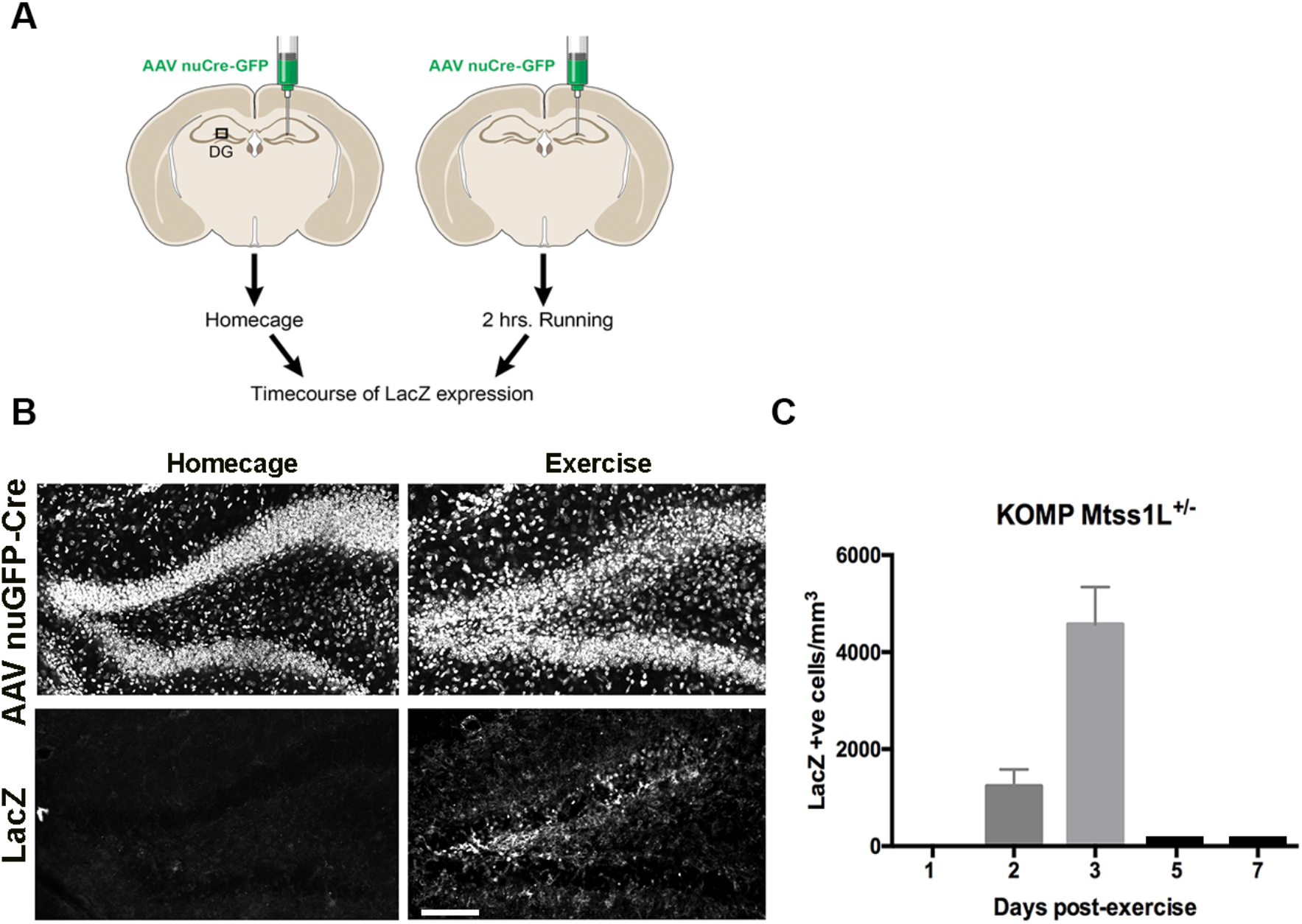
Timecourse of LacZ expression in Mtss1L reporter mice. **(A)** Schematic representation of the experimental paradigm used for the Mtss1L reporter mice. AAV9-Cre-nuGFP viral vector was injected unilaterally into the dentate gyrus of KOMP Mtss1L+/− mice in order to generate the LacZ reporter allele. Two weeks after stereotaxic injections, KOMP Mtss1l+/− were either housed in their homecage or exposed to 2 hrs of voluntary exercise. Brain sections were processed for LacZ immunohistochemistry at several time points. **(B)** Representative images of dentate gyrus show robust AAV9-Cre-nuGFP expression in granule cells (top panels) in both control (homecage) and exercised mice. LacZ was detected only in dentate granule cells of mice at 3 days post-exercise (bottom, right) whereas no expression was observed in homecage littermates (bottom, left). Scale bar: 250 μm. Likewise, no expression was detected in the non-virally injected hemisphere (data not shown). **(C)** LacZ expression in granule cells peaked at 3 days post-exercise and was not detectable at 5 or 7 days post-exercise (LacZ+ cells/mm^3^ 2 days post-exercise: 1235±345, n=3; 3-days post-exercise: 4573±767,n=3, one-way ANOVA p<0.0001).

Mtss1L belongs to the inverse-BAR (Bin, Amphiphysin and Rvs, I-BAR) protein family, a class of proteins that function by promoting curvature in membranes. BAR domain proteins such as amphyphysin promote convex curvatures including invaginations that are involved in endocytosis or synaptic vesicles at presynaptic nerve terminals *(22).* In contrast, I-BAR proteins promote concave structures typically seen in outward-projecting membrane protrusions. Thus, we hypothesized that Mtss1L expression mediated exercise-dependent formation or rearrangement of synapses in postsynaptic dendrites.

As proof of principle, we first examined the localization of endogenous Mtss1L in hippocampal neurons *in vitro* following exposure to brain-derived neurotrophic factor (BDNF), which is upregulated by exercise and involved in activity-dependent synaptic plasticity *(23, 24)*. Mtss1L immunoreactivity was detectable only after BDNF treatment (Fig. 5A), consistent with the activity-dependent expression pattern we observed *in vivo.* Mtss1L colocalized with the somatodendritic marker MAP2 along dendrites and in dendritic spines (Fig. 5A). To determine whether Mtss1L could induce spine-like protrusions, we compared spine density in cultured hippocampal neurons transfected with mCherry or mCherry-Mtss1L. At DIV 14, Mtss1L overexpression markedly increased spine-like protrusions (Fig. 5B) including filopodia and mushroom spine subtypes (fig. S6). The somatodendritic pattern of overexpressed mCherry-Mtss1L overlapped the endogenous Mtss1L expression (Fig. 5B, yellow). mCherry-Mtss1L-expressing dendritic spines also were labeled with co-transfected PSD-95-FingR-GFP, (Fibronectin Intrabody generated with mRNA display) that binds to endogenous PSD-95 *(25)*. PSD-95 intrabody labeling of excitatory synapses in living neurons indicated that the protrusions induced by Mtss1L also contain postsynaptic proteins (Fig. 5C). *In vivo,* overexpression of mCherry-Mtss1lL by DNA electroporation at P0 increased spine density of mature dentate granule cells at P21 (Fig. 5D). These results indicate that Mtss1L is expressed in dendrites and can induce spine-like protrusions, consistent with a putative role in synaptic plasticity.

**Figure 5:**
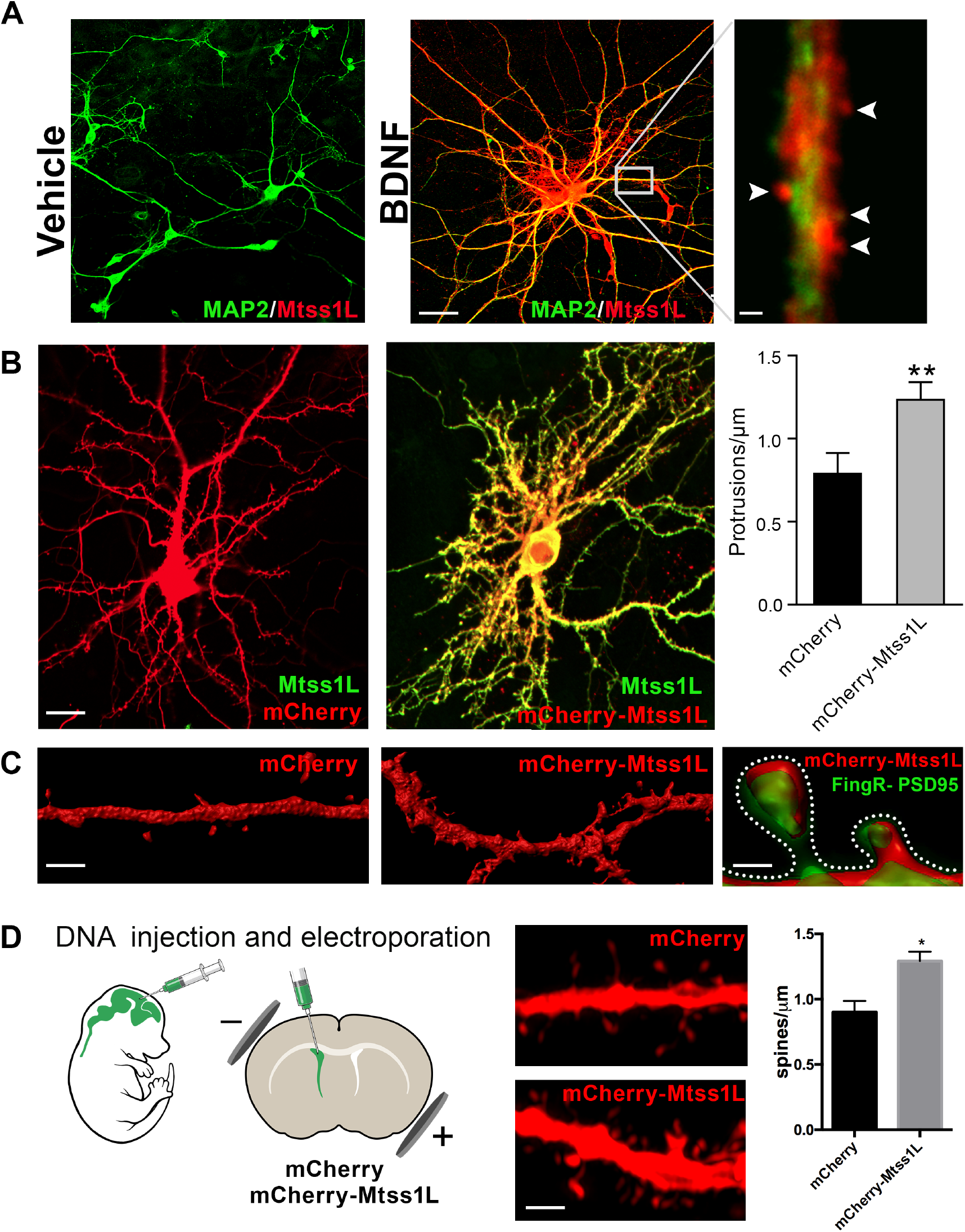
Induced Mtss1L expression *in vitro* and *in vivo.* **(A)** Primary hippocampal neurons were cultured with or without BDNF from DIV 7-14 to examine activity-dependence of Mtss1L expression. Representative images of hippocampal neurons co-stained with anti-Mtss1L (red) and the somatodendritic marker anti-MAP2 (green) showed robust Mtss1L immunoreactivity in BDNF-treated cells (middle), but not in the vehicle-treated controls (left). Scale bar: 20μm. High magnification image of a single dendrite in a BDNF-treated neuron showed localization of Mtss1L (red) in dendritic shaft and spines (right, arrowheads, scale bar: 1.5μm). **(B)** Representative images of mouse hippocampal neurons transfected with mCherry or Mtss1L-mCherry and stained with anti-Mtss1L (green). Scale bar: 12μm. Ectopic expression of Mtss1L markedly increased the number of dendritic protrusions (mCherry: 0.9±0.09, Mtss1L-mCherry: 1.3±0.07, unpaired t-test, p= 0.004, n=3). **(C)** Higher magnification images of dendritic segments from hippocampal neurons transfected with mCherry (left) or Mtss1L-mCherry (middle) shows the increased number of protrusions. Scale bar: 3μm. Merged image at right of PSD-95.FingR-GFP and Mtss1L-mCherry in co-transfected neurons demonstrates that the protrusions contained postsynaptic proteins. Scale bar: 0.5 μm. **(D)** DNA solution was injected into the lateral ventricle of P0 pups followed by gene delivery by electroporation. Representative dendritic segments of dentate granule cells expressing control plasmid mCherry (top) or mCherry-Mtss1L (bottom) 21 days postelectroporation. Scale bar: 3μm. Mtss1L-mCherry expressing cells show increased dendritic protrusions *in vivo* (mCherry, 0.9±0.03, n=3, Mtss1L-mCherry, 1.3±0.07, n=3, unpaired t-test, p=0.004).

To determine whether Mtss1L was responsible for the increase in synapses in exercise TRAPed cells, we used a lentivirus expressing shRNAs targeting Mtss1L. Validating the efficacy of the shRNAs, shMtss1L reduced mCherry-Mtss1L mRNA and protein expression in HEK293T cells and in primary hippocampal neurons treated with BDNF (fig. S7). To knockdown Mtss1L *in vivo* (Fig. 6A), lentiviral shMtss1L-GFP or shscramble-GFP were injected into the dentate gyrus of Fos-TRAP mice. Three weeks post-viral injection, we assessed spine density and excitatory synaptic function in TdT^+^ (exercise-TRAPed) granule cells that also co-expressed GFP as a marker for shRNA expression. At 3 days post-exercise, knockdown of Mtss1L occluded the exercise-induced increase in dendritic spines in the OML (Fig. 6B middle panel top, and right panel). The shscramble-GFP, injected into the contralateral dentate gyrus of the same mice, had no effect (Fig. 6B right panel). Viral knockdown of Mtss1L in exercise-TRAPed cells also prevented the increase in evoked EPSCs in simultaneous paired recordings from TdT^−^ and TdT^+^/GFP+ granule cells (Fig. 6C-D). The amplitude of EPSCs following viral knockdown in exercise-TRAPed cells was the same as neighboring cells that were not activated by exercise (Fig 6D, right panel). Intrinsic membrane properties were not affected by Mtss1L knockdown (Table S4). Figure 6E includes the results from Figure 2 (“exercise-TRAPed OML” (red) and “exercise-TRAPED MML” (black), compared to the effects of Mtss1L shRNA (yellow), demonstrating that knockdown of Mtss1L was necessary to explain the exercise-induced increase in excitatory synapses. Importantly, evoked EPSCs in granule cells that only expressed GFP+shRNA were unaffected, demonstrating that the effect of Mtss1L was dependent on activity-dependent expression (fig. S8).

**Figure 6:**
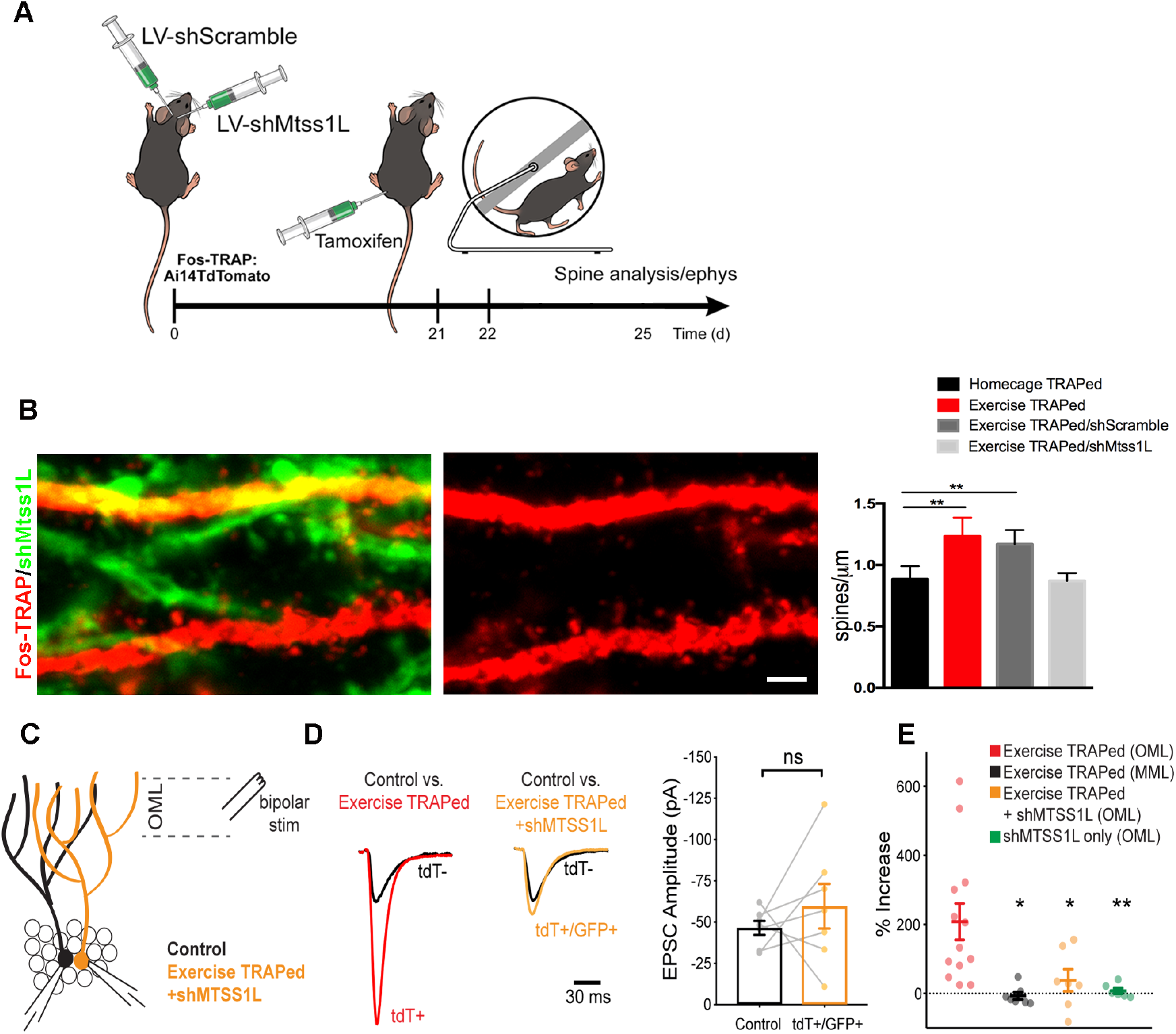
Mtss1L knockdown in exercise-TRAPed cells blocked the exercise-induced increase in dendritic spines and synaptic activity at 3 days post-exercise. **(A)** Effects of Mtss1L knockdown on the spine density of exercise-TRAPed cells were assessed by stereotaxically injecting a control shScramble-GFP lentivirus into the left hemisphere and a shMtss1L-GFP lentivirus in the right hemisphere of Fos-TRAP mice. After 21 days, dentate granule cells were exercise-TRAPed and analyzed 3 days later as shown in the schematic. **(B)** The left panel shows representative OML dendritic segments of exercise-TRAPed cells (red), exercise-TRAPed cells co-expressing shRNA (orange) next to several dendrites expressing only shRNA (green). The middle panel shows the dendritic field isolated in the red channel to allow comparison of dendritic spines in exercise-TRAPed cells at 3 days post exercise with (top, middle panel) and without (bottom, middle panel) co-expression of the Mtss1L shRNA. Mtss1L shRNA, but not shScramble, blocked the expected increase in dendritic spines in exercise-TRAPed cells. Summary graph at right shows dendritic spine density in the dentate OML for each condition (Homecage TRAPed: 0.8±0.1, n=4, Exercise TRAPed: 1.2±0.2, n=5, Exercise TRAPed/shScramble: 1.17±0.1, n=4, Exercise-TRAPed/shMtss1L: 0.87±0.06, n=5, One-way ANOVA, multiple comparisons Dunnett’s test, p<0.01). **(C)** Schematic of simultaneous whole-cell recordings from control (black) and exercise TRAPed cells co-expressing shRNA-Mtss1L (orange) to assess functional synaptic activity. Lateral perforant path axons were stimulated in the OML. **(D)** Superimposed EPSCs from a representative control cell (black) and a simultaneously recorded exercise-TRAPed cell (red) showed a large increase in amplitude in the exercise-TRAPed cells, as quantified in Fig. 1E. In contrast, superimposed EPSCs from a control cell (black, 46.5 ± 4.2 pA) and an exercise-TRAPed cell co-expressing shMtss1L (orange, −59.6 ± 13.5 pA) showed no increase in amplitude (p = 0.42, paired t-test, 7 cell pairs from 5 mice). Traces are normalized and scaled relative to control cells (black) for presentation. **(E)** Summary plot across experimental conditions. The exercise-TRAPed increase in EPSC amplitude in the OML was blocked by Mtss1L shRNA expression in exercise-TRAPed cells. (Percent increase in EPSC amplitudes: exercise-TRAPed – OML stimulation, 208.2 ± 52.8 %, n = 13 cells, 8 mice; exercise-TRAPed – MML stimulation, – 6.7 ± 10.1 %, n = 7 cells, 4 mice; exercise TRAPed + shMtss1L – OML stimulation, 38.4 ± 32.1 %, n = 7 cells, 5 mice; shMtss1L only – OML stimulation, 7.8 ± 8.0 %, n = 6 cells, 3 mice, p = 0.002, 1-way ANOVA; exercise-TRAPed shMtss1L, p = 0.025; shMtss1L only, p = 0.011; exercise-TRAPed MML, p = 0.004; Dunnett’s test).

## Discussion

Our experiments were designed to test the cellular and molecular response to acute exercise with an emphasis on time periods during which synapses might form or reorganize. This approach differs from studies of sustained or chronic exercise that likely involve systemic as well as neural mechanisms *(3, 4, 26).* Such studies have shown effects of exercise on many aspects of hippocampal function, including neurogenesis, synaptic plasticity, structural changes at synapses and dendritic spines as well as behavioral effects on learning and memory *(1, 2)*. Our goal was to examine the transcriptional response in the days following acute exercise, which led to the identification of a novel effector of structural plasticity, Mtss1L.

Although the Fos-TRAP method is limited by the time required for reporter expression (ca. 1 day post-exercise), the immediate early gene Egr1 was upregulated (Fig. 3C), indicating that our protocol detected activity-dependent gene expression in mature granule cells. Although the presence of the running wheel itself could act as a novel object, there was only a small increase in Fos-TRAPed cells when the wheel was fixed to prevent running (data not shown). It is interesting that adult-generated granule cells in the subgranular zone were not activated by two hours of exercise (data not shown). Although it is well known that chronic exercise increases adult neurogenesis *(27)*, progenitors and newborn neurons do not receive direct excitatory perforant path input for several weeks *(28)*, perhaps explaining their lack of activation in our experiments.

Our results provide the first evidence for activity-dependent expression of an I-BAR protein, and suggest that I-BAR proteins can affect both constitutive dendritic spine formation *(29)* as well as experience-dependent remodeling of synapses. The immediate family members of Mtss1L include MIM/Mtss1 and IRSp53 *(30)*, but their function in the nervous system is just beginning to be explored. For example, Mtss1L expression was previously detected in radial glial cells during development as important for membrane protrusions and end-feet *(20)*. The related I-BAR protein, MIM/Mtss1, is involved in cerebellar synapse formation with signaling including PIP2-dependent membrane curvature and subsequent Arp2/3-mediated action polymerization *(29)*. MIM and another member of the I-BAR subfamily, IRSp53, have also been implicated in aspects of synapse formation during development *(31, 32).*

At a circuit level, the laminar-specific increase in synaptic function suggests that acute exercise had a network-specific effect. Entorhinal inputs to the outer molecular layer, the distal dendrites of mature granule cells, have been previously associated with contextual information and particularly the “how” and “when” aspects of encoding memories *(33)*. How might these results related to the beneficial effects of exercise? One possibility consistent with our data is that exercise acts as a preconditioning signal that primes exercise-activated neurons for contextual information incoming during the several days following exercise. This represents a broader time window than is usually associated with short-term plasticity. For example, human studies give support for the idea that exercise within 4 hours of a learning task improved memory performance *(34)*. It will be interesting to examine if exercise enhances the pattern of granule cell responses to spatial or context-specific tasks. Our identification of Mtss1L as an activity-dependent I-BAR domain protein makes it ideally suited to act at an early mediator of structural plasticity following neural activity.

## Materials and Methods

### Mice

All procedures were performed according to the National Institutes of Health Guidelines for the Care and Use of Laboratory Animals and were in compliance with approved IACUC protocols at Oregon Health & Science University. Both female and male mice from a C57BL6/J background were used for experiments, aged 6-8 weeks at the time of surgery. TRAP mice were heterozygous for the *Fos-^iCreER^* allele, with some also heterozygous for the *B6;129S6-Gt(ROSA)26Sortm14(CAG-tdTomato)/Hze/ J*(*Ai14;*JAX#007908) allele for experiments involving tdTomato labeling. Cryopreserved Mtss1l^tm1a(KOMP)Wtsi^ sperm was obtained from the Knockout Mouse Project (KOMP) Repository (The Knockout Mouse Project, Mouse Biology Program, University of California, Davis, CA, USA)under the identifier CSD79567. The Mtss1l^tm1a(KOMP)Wtsi^ mouse line was re-derived by *in vitro* fertilization at OHSU transgenic core. Genotype was confirmed by PCR. All mice were housed in plastic cages with disposable bedding on a standard light cycle with food and water available ad libitum.

### Stereotaxic Injections

Mice were anesthetized using an isoflurane delivery system (Veterinary Anesthesia Systems Co.) by spontaneous respiration, and placed in a Kopf stereotaxic frame. Skin was cleaned with antiseptic and topical lidocaine was applied before an incision was made. Burr holes were placed at x: ±1.1 mm, y: −1.9 mm from bregma. Using a 10 μl Hamilton syringe with a 30 ga needle and the Quintessential Stereotaxic Injector (Stoelting), 2 μl mixed viral stock (1 μl of each virus) was delivered at 0.25 μl/min at z-depths of 2.5 and 2.3 mm. The syringe was left in place for 1 min after each injection before slow withdrawal. The skin above the injection site was closed using veterinary glue. Animals received post-operative analgesia with topical lidocaine and flavored acetaminophen in the drinking water.

### TRAP induction

Tamoxifen was dissolved at 20 mg/ml in corn oil by sonication at 37°C for 5 min. The dissolved tamoxifen was then stored in aliquots at −20°C for up to several weeks or used immediately. The dissolved solution was injected intraperitoneally at 150 mg/kg. For exercise-TRAP, tamoxifen was administered 23 hours before exposure to the running wheel. Experiments were begun 48 hours after tamoxifen administration to allow for reporter expression.

### Voluntary Exercise Protocol

Both homecage and exercise groups were housed together until 7 days before the experiment, when they were singled housed in an oversized (rat) sedentary cage (43×21.5 cm^2^) to allow acclimation in the novel environment before tamoxifen administration. At 23hrs post-tamoxifen injection, a running wheel was introduced in the cage of animals in the exercise group and mice had free access to running for 2hrs at the beginning of the dark period, after which the running wheel was removed from the cage. Total distance (km) was measured using an odometer. All mouse groups were handled for 5 days before tamoxifen administration.

### Immunohistochemistry

Mice were terminally anesthetized, transcardially perfused with saline and 20ml of 4% paraformaldehyde (PFA), and brains were post-fixed overnight. Coronal sections (100μm) of the hippocampus were collected and permeabilized in 0.4% Triton in PBS (PBST) for 30 min. Sections were then blocked for 30 min with 5% horse serum in PBST and incubated overnight (4°C) with primary antibody in 5% horse serum/PBST. After extensive washing, sections were incubated with the appropriate secondary antibody conjugated with Alexa 488, 568 or 647 (Molecular Probes), for 2 hours at room temperature. They were then washed in PBST (2 x 10 min) and mounted with Dapi Fluoromount-G (SouthernBiotech). The primary antibodies used were: anti-cfos (1:500, Santa-Cruz), anti-tdTomato (1:500, Clontech), anti-MAP2 (1:500, Sigma), anti-ABBA/Mtss1L (1:500, Millipore), anti-VGLUT2 (1:500, Synaptic Systems). All antibodies have been well characterized in prior studies in our laboratory, and staining was not observed when the primary antibody was omitted.

### Imaging and Morphological Analysis

For imaging we used an LSM 7 MP laser scanning microscope (Carl Zeiss MicroImaging; Thornwood, NY). All cell and dendritic spine counting and dendritic arbor tracing were done manually with ImageJ and Imaris (National Institutes of Health; Bethesda, MD). For quantification of immunopositive cells, six hippocampal slices at set intervals from dorsal to ventral were stained per animal; 3-6 animals were analyzed per group. A 49 μm z-stack (consisting of 7 optical sections of 7μm thickness) was obtained from every slice. Positive cells were counted per field from every z-stack, averaged per mouse and the results were pooled to generate group mean values. For analysis of spine density of dentate granule cells, sections were imaged with a 63x objective with 2.5x optical zoom. The span of z-stack was tailored to the thickness of a single segment of dendrite with 0.5μm distance between planes. Dendritic spines were imaged and analyzed in the middle (MML) and outer molecular (OML) layers of the dentate gyrus. The MML and OML were distinguished based on the pattern of VGluT2 immunofluorescence, which begins at the border between the inner molecular layer and middle molecular layer. MML dendritic segments were therefore imaged at the beginning of the VGluT2 staining nearest the granule cell body layer, whereas OML dendritic segments were imaged at the distal tip of the molecular layer. The same microscope settings (laser intensity and gain) were used for each experimental group analyzed. Slides were coded and imaged by an investigator blinded to experimental condition.

### Laser Capture Microdissection (LCM)

To obtain nuclei of exercise-TRAPed cells, fresh frozen coronal brain sections (12 μm) were collected by cryostat sectioning on polyethylene napthalate membrane slides. Sections were fixed for 1 min in 75% ethanol, followed by a final wash in 100% ethanol for 1 min. Pools of 150 RFP^+^ and RFP^−^ individual cells were laser captured from the granular cell layer of the dentate gyrus of each mouse using a Leica Microsystem LMD system under a HC PL FL L 63/0.60 XT objective and collected in lysis buffer (50 ul RLT buffer, Qiagen). Total RNA from LCM samples was isolated separately from each animal.

### RNA extraction and RNAseq library preparation

RNA was extracted from laser-captured materials using RNeasy mini-elute column with a modified protocol. Briefly, cells were picked from laser capture microscopy onto caps of 0.5ml Eppendorf tubes, 50ul RLT buffer was added immediately, and placed on ice. After spinning down the samples, 300ul RLT was added to each sample, mixed well, and vortexed for 30s. After brief spin, 2ng tRNA was added as carrier and 625μl 100% ETOH was added into each sample and mixed well. After a quick spin, samples were loaded onto RNeasy mini-elute columns, washed, dried and eluted in 10ul RNAse-free water. 8ul RNA was used for reverse transcription using Superscribe reverse transcriptase 2ul, DTT 0.1M 0.5ul, 5x RT buffer 4ul at 42degrees Celsius for 1.5hr then heat inactivated at 75 °C for 20 minutes (self-developed protocol for RNA extraction, superscribe reverse transcriptase was from Clontech, protocol adopted from manufacturer’s protocol for smarter single cell RNAseq v4). TSO (AAGCAGTGGTATCAACGCAGAGTACrG+GrG was ordered from Exicon. After RNA, PCR was carried out using Seqamp DNA polymerase and amplification PCR primer (AAGCAGTGGTATCAACGCAGAGTAC). Half of amplified cDNA was used for qPCR and the other half made into sequencing libraries using Nextera XT DNA library prep kit (Illumina) Quality of cDNA libraries were checked on Bioanalyzer using DNA high sensitivity Chip. 2nM library dilutions were mixed and denatured, 1.8pmol mixed library was sequenced on Nextseq500 (Illumina).

### RNAseq analysis

After sequencing, sequencing quality was checked by fastqc, raw reads were aligned with STAR 2.5b on Illumina Basespace. Raw reads were then plotted as a violin plot using the R ggplot2 package. Genes with reads lower than 64 in all samples were filtered away and dataframe was replotted to check data distribution. Individual samples that differed significantly in their count distribution due to technical issues were discarded from further analysis. Data dispersion and differential expression analysis was done on filtered reads using DEseq2 unpaired analysis. FDR<0.15 fold enrichment > 1.5 fold were called as enriched genes were plotted in MA-plot using R script. Genes enriched in TRAPed cells at 3 day post exercise were further analyzed by PantherGO for GO terms enrichment (release 20160321, GO Ontology database Released 2016-05-20); protein-protein reactions analyzed by Cytoscape application Biogenet.

### Real-time qPCR

To quantify gene expression levels, amplified cDNA was mixed with the iQ5 SYBR Green PCR master mix (Biorad) and 5 pmol of both forward and reverse primers were used. β-actin was amplified as an internal control. For qPCR of miRNAs, miRNA was converted to cDNA using the QuantiMiR reverse transcription kit (System Biosciences, Mountain View, CA). Briefly, RNA was polyadenylated with ATP by poly(A) polymerase at 37°C for 1 hr and reverse transcribed using 0.5 μg of poly(T) adapter primer. Each miRNA was detected by the mature DNA sequence as the forward primer and a 3’ universal reverse primer provided in the QuantiMir RT kit. Human small nuclear U6 RNA was amplified as an internal control. qPCR was performed using Power SYBR Green PCR Master Mix (Applied Biosystems). All qPCR performed using SYBR Green was conducted at 50°C for 2 min, 95°C for 10 min, and then 40 cycles of 95°C for 15 s and 60°C for 1 min. The specificity of the reaction was verified by melt curve analysis. Relative expression was calculated using mouse Ubc as internal control. A table of primers used is included, Table S4.

### Primary neuronal hippocampal cultures and *in vitro* transfections

Hippocampi from E18-19 embryos were isolated as described previously (35) and placed on ice. The tissue was chopped into 1 mm cubes with fine scissors and transferred to a 15 ml tube using a Pasteur pipette. This tube was placed in a water bath at 37 °C with the papain solution, separately, and both were allowed to equilibrate at this temperature for 5 min. The tissue was then carefully transferred into the papain solution and placed on an orbital shaker at 100 rpm at 37 °C for 15 min. The tissue was then gently transferred from the papain solution into a 15 ml tube containing 2 ml HBSS. This step was repeated 2X to remove any residual papain. The tissue was then triturated using fire polished Pasteur pipettes. Following trituration and centrifugation, the single cell suspension was seeded on poly-D-Lysine coated coverslips in 24-well plates at 10^4^ cells/well in Neurobasal Plating Media with B27 Supplement [1 ml / 50 ml], 0.5 mM Glutamine Solution, Penicillin (10,000 units / ml)/Streptomycin (10,000 μg / ml) [250 μl / 50 ml], 1mM HEPES (MW 238.3 g / mol), 10% Heat-Inactivated horse serum. HI-horse serum was removed by half-changes of the plating media beginning at 24 hours.

For transfection of cultured neurons, DNA-Lipofectamine 2000 complexes were prepared as follows: 1μg of DNA was diluted in 50μl of Neurobasal medium and 2 μl of Lipofectamine 2000 was diluted in another 50μl of Neurobasal medium. Five minutes after dilution of Lipofectamine 2000, the diluted DNA was combined with the diluted lipid. The solutions were gently mixed and then left for 20 min at room temperature to allow the DNA–Lipofectamine 2000 lipoplexes to form. 100 μl of transfection complex was added to each well containing cells and medium and the cells were incubated at 37 °C in a humidified incubator with 5% CO_2_ for 1hr after which transfection media was changed. Transfection efficiency was assayed 48hrs posttransfection by fluorescence microscopy.

### *In vivo* electroporation into postnatal dentate granule cells of the dentate gyrus

Postnatal day 0 (P0) electroporation of the dentate gyrus was performed accordingly to a previously published method *(36)*. In brief, mice were anesthetized by hypothermia, and then a sharp glass electrode with a beveled tip containing plasmid (1 μg/μl in TE mixed with 0.05% Fast Green) was inserted through skin and skull. One μl of DNA solution was injected into the lateral ventricle (LV). Correct injection was confirmed by trans-cranial visualization of Fast Green in the LV in the dissecting microscope. Successfully-injected pups were immediately electroporated with tweezer electrodes (5 mm platinum) positioned onto the brains as described by Ito et al. *(36)*. Five pulses of 100 V and of 50 ms duration were given at 950 ms intervals. Electroporated animals were placed in a recovery chamber at 37 °C for several minutes and then returned to their mother.

### Plasmid construction

PLVX-cherry-C1 plasmid was from Clontech and Mtss1L coding region was cloned into the C-terminus of mCherry between BsrG1 and SmaI sites to generate fusion construct of PLVX-cherryC1-Mtss1L (Mtss1L coding region sequence Accession number B), FUCW-T2A-Mtss1L/ABBA was constructed using NEB HiFi DNA builder assembly kit. shABBA constructs were either obtained from Dharmacon or designed by Biosettia and subcloned into PLV-UBC-GFP-mU6-shRNA using NEB HIFI DNA assembly mix. The folllowing shRNA sequences were used: Mtss1L1:AAAAGGCCGTTTCTGCACCTTTATTGGATCCAATAAAGGTGCAGAAA CGGCC, Mtss1L2: TCTCTACTAGGCTGTGCCT, Scramble:AAAAGCTACACTATCGAGCAATTTTGGATCCAAAATTGCTCGATAGTGTAGC

### Slice Physiology

Acute brain slices were prepared as previously described *(37)*. In summary, 8-week-old male and female mice were anesthetized with 4% isoflurane and injections of 1.2% avertin (Sigma-Aldrich). Mice were perfused with 10 mL of ice-cold cold N-methyl-D-glucamine (NMDG)-based cutting solution, containing the following (in mM): 93 NMDG, 30 NaHCO3, 24 glucose, 20 HEPES, 5 Na-ascorbate, 5 N-acetyl cysteine, 3 Na-pyruvate, 2.5 KCl, 2 thiourea, 1.2 NaH2PO4, 10 MgSO4, and 0.5 CaCl2. Transverse 300 μm thick hippocampal sections were cut in ice-cold NMDG solution on a Leica VT1200S vibratome. Slices recovered in warm (34 C) NMDG solution for 15 minutes. Slices were transferred to room temperature in standard ACSF and allowed to recover for at least 1 hour prior to recording.

Dentate granule cell whole-cell recordings were made with 3-5 MΩ borosilicate glass pipets filled with a Cs^+^-based internal solution containing (in mM): 113 Cs-gluconate, 17.5 CsCl, 10 HEPES, 10 EGTA, 8 NaCl, 2 Mg-ATP, 0.3 Na-GTP, 0.05 Alexa Fluor 488, pH adjusted to 7.3 with CsOH. Osmolarity was adjusted to 295 mOsm and QX-314-Cl (5 mM; Tocris Bioscience) was included to block unclamped action potentials. The junction potential was 8 mV and left uncorrected. Granule cells were visually identified on an Olympus BX-51WI microscope using infrared differential interference contrast imaging. To ensure relatively equal stimulation intensity between cells during paired simultaneous tdT+ and tdT-recordings, neighboring tdT+ and tdT-(control) cells were selected. Cells were only considered “neighbors” if they were within roughly ±2 cell bodies distance apart in the x and y planes, and roughly ±1 cell body distance in the z plane. Granule cells were filled with AlexaFluor 488 to ensure dendritic overlap. Bi-polar stimulating electrodes (FHC) were placed in distally in the OML and medially in the MML, to ensure OML and MML specific stimulation of lateral and medial perforant path, respectively *(38)*. Electrical stimulation was delivered with a constant current stimulator (Digitimer, Inc); intensity was tittered to produce 50 – 100 pA EPSCs in tdT-cells. Proximity to the stimulating electrode and recording configuration was randomly assigned. Cells were voltage-clamped to −70 mV and signals were amplified using two AxoClamp 200B amplifiers (Molecular Devices), Bessel filtered at 5 kHz, and captured at 10 kHz with an analog-to-digital converter (National Instruments). Data were collected and analyzed in Igor Pro 8 (Wavemetrics) using a custom script. Averages were comprised by the mean of 15 consecutive sweeps.

### Statistical analysis

Sample sizes were based on pilot experiments with an effect size of 20% and a power of 0.8. Littermates were randomly assigned to control or exercise groups. Criteria were established in advance based on pilot studies for issues including data inclusion, outliers, and selection of endpoints. Criteria for excluding animals from analysis are listed in the methods. Mean ± SD was used to report statistics for all apart from electrophysiology experiments where Mean ± SE was used. The choice of statistical test, test for normality, definition of N, and multiple hypothesis correction where appropriate are described in the figure legends. Unless otherwise stated, all statistical tests were two-sided. Significance was defined as p =0.05. All statistical analyses were performed in Prism or Igor Pro 8.

## Acknowledgements

We thank the following for assistance with our experiments: Stefanie Kaech Petrie, Aurelie Snyder, Crystal Shaw and the Advanced Light Microscopy Core (P30 NS061800); Shannon McWeeney and Sophia Jeng in the OHSU Bioinformatics core; Laura Villasana and Jocelyn Santiago-Perez (behavioral testing); Francesca Cargnin (electroporation); Lev Federov and OHSU transgenic core (generation of KOMP mice). This work was supported by the Ellison Foundation and NIH NS080979 (RHG and GLW), Department of Veterans Affairs Merit Review Award I01-BX002949 (ES); a Department of Defense CDMRP Award W81XWH-18-1-0598 (ES); a National Institutes of Health (NIH) Grant F31-NS098597 (WDH); and a fellowship from Ronni Lacroute (CC). The contents of this manuscript do not represent the views of the U.S. Department of Veterans Affairs or the United States government. RNA seq data generated in the manuscript have been deposited at: https://www.ncbi.nlm.nih.gov/bioproject/PRJNA481775

## Supplementary Materials for

**Fig. S1:**
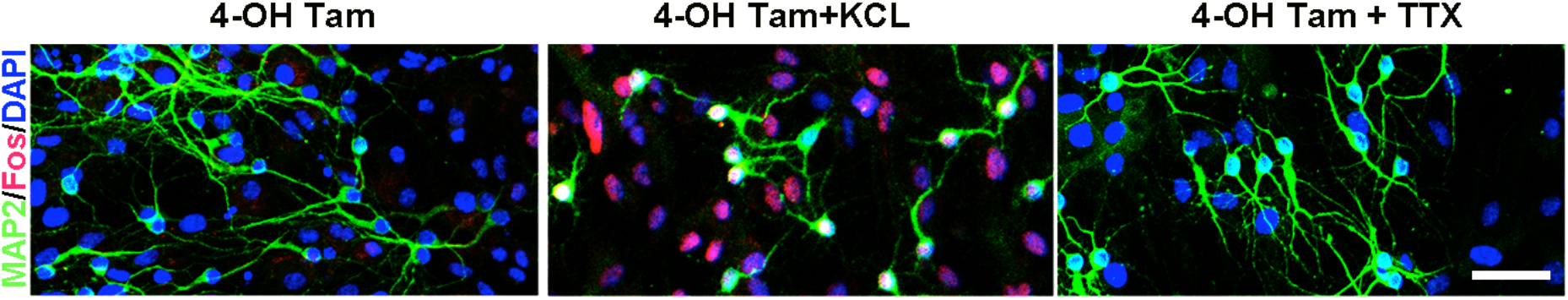
*In vitro* validation of Fos-TRAP method. To validate the sensitivity of the Fos-TRAP method, primary hippocampal cultures were derived from 2-day old Fos-TRAP pups, transduced with an AAV-flex-H2B-mRFP virus, and then 5 days later treated with 4-OH tamoxifen (4-OH Tam, 1μM) for 48 hrs. As expected, H2B-mRFP labelling was sparse in these low density cultures (left panel), whereas KCl-induced depolarization (20mM, 30 min) resulted in a robust increase in Fos:H2B-mRFP expression (red, middle panel). Inhibition of neuronal activity in the cultures for 48 hours with TTX (500nM), NBQX (2.5μM) and CPP (5μM) completely abolished Fos-TRAP: H2B-mRFP expression (right panel). Scale bar: 40 μm.

**Fig. S2.**
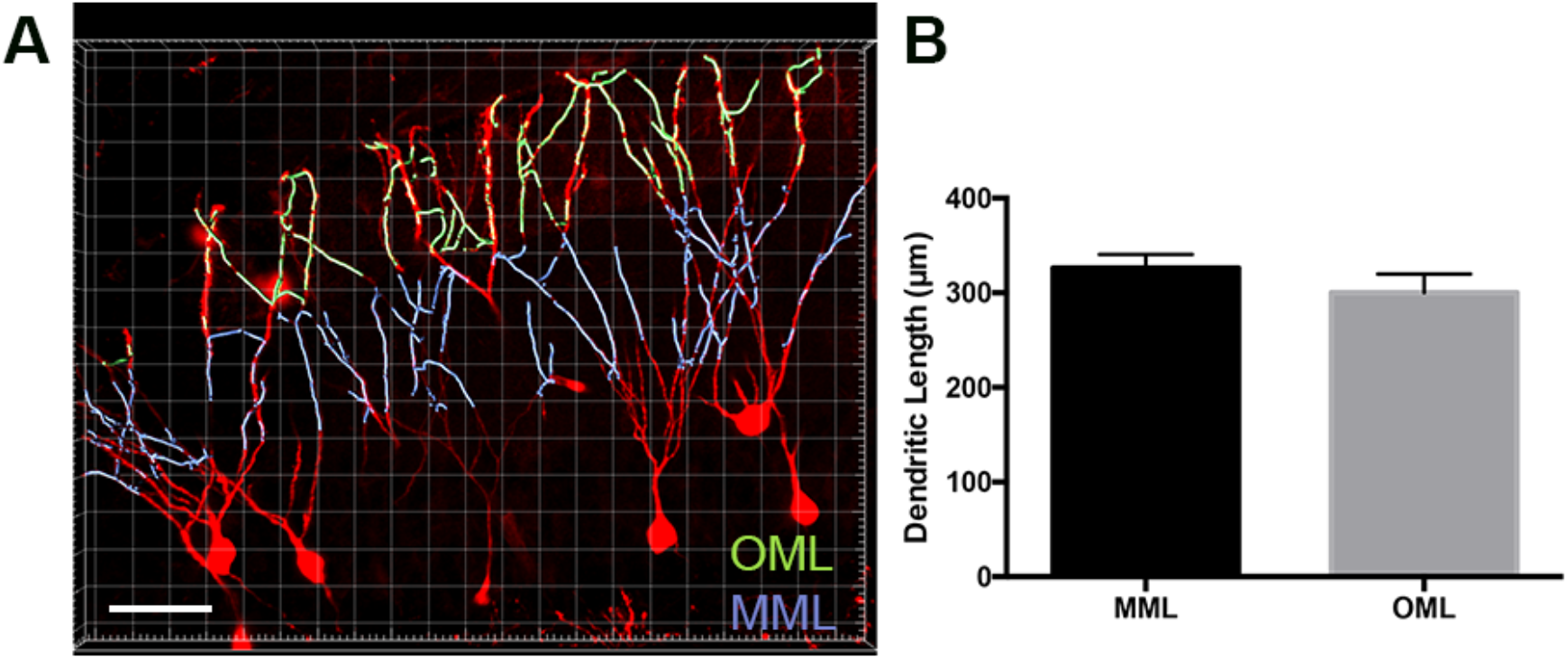
Dendritic lengths were equal in outer and middle molecular layers of exercise TRAPed cells. **(A)** Representative image of dendrites of the outer molecular layer (light green) and middle molecular layer (light purple) of exercise-TRAPed cells (red) 3 days post-exercise. The band of VGluT2 immunofluorescence labeled only the OML and MML, with the outer half representing OML and the inner half representing MML for quantification. Scale bar: 40 μm. **(B)** There was no difference in dendritic lengths per cell between MML and OML cells (MML: 326±14 μm, n=3, OML: 300±20, n=3, unpaired t-test, p=0.14, 49 μm stack). Masking was used to better visualize the OML and MML in the image.

**Fig. S3.**
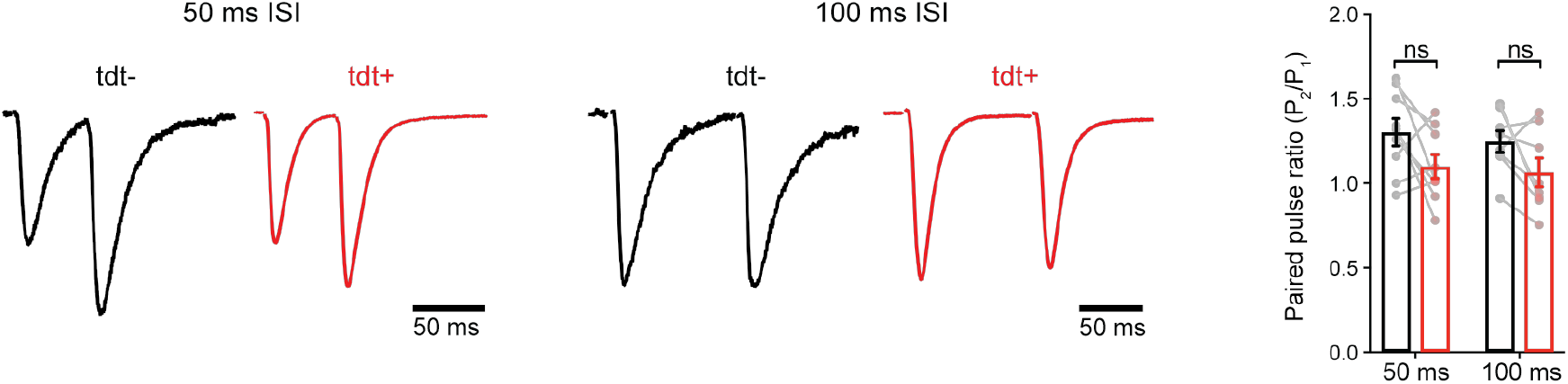
Paired-pulse ratio (PPR) in control and exercise-TRAPed granule cell paired recordings. **(A)** Representative EPSCs from paired-pulse experiments during 50 ms interstimulus interval (ISI) OML stimulation. Responses are scaled on the y-axis relative to the first peak (PPR, 50 ms ISI: tdT-, 1.30 ± 0.08, tdT^+^, 1.10 ± 0.07, n = 9 cells, p = 0.0770, paired t-test). Stimulus artifacts were blanked for presentation. **(B)** Same as in A, but with 100 ms ISI PPR, 100 ms ISI: tdT-, 1.25 ± 0.06, tdT^+^, 1.06 ± 0.08, n = 8 cells, p = 0.09, paired t-test) **(C)** Quantification and summary of paired-pulse experiments. There was no change in PPR at 50 ms ISI (p = 0.08, paired t-test) or 100 ms ISI (p = 0.09, paired t-test). ns, not significant.

**Fig. S4.**
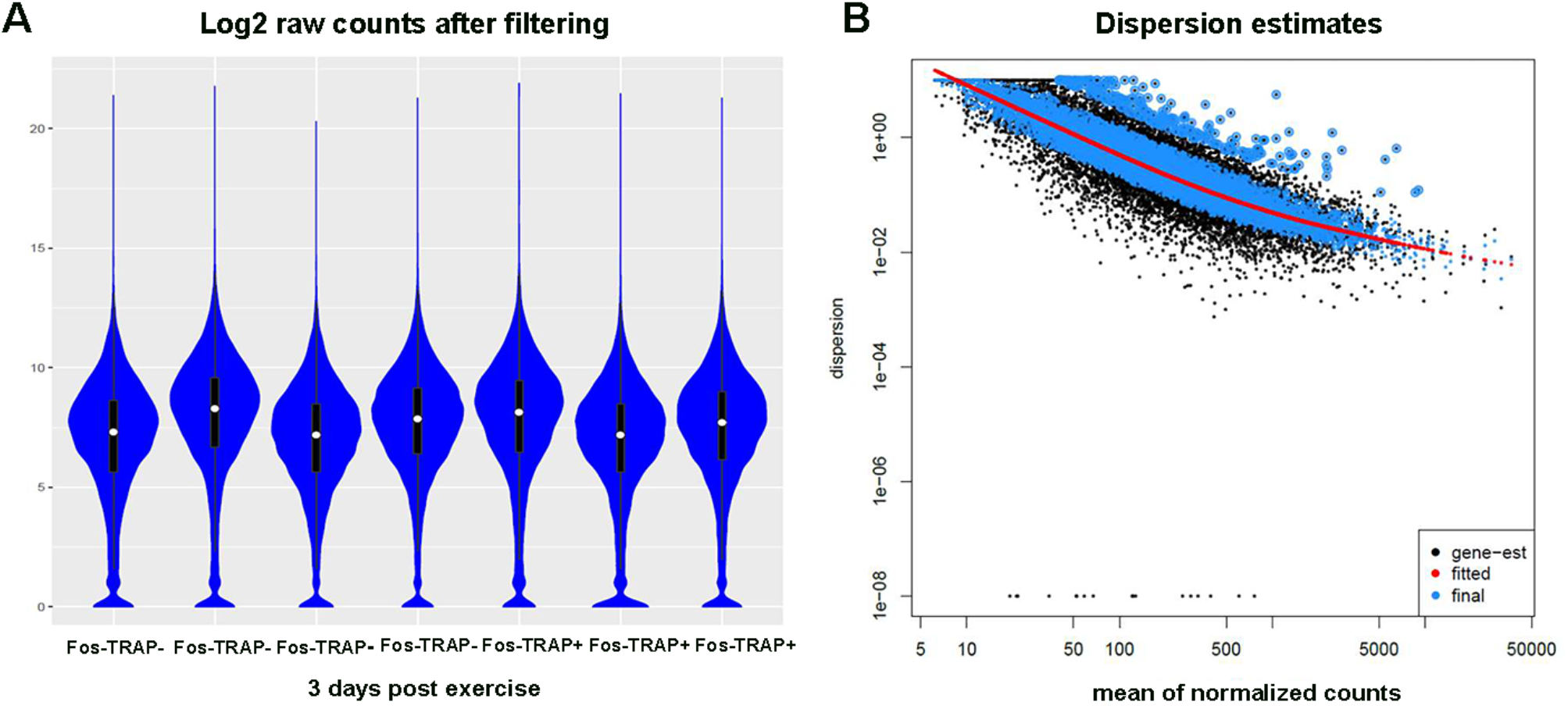
Filtering, and dispersion estimation of sequencing data used for transcriptomics. **(A)** Log2 raw counts dataframe after filtering transcripts with rowmax <64 counts are displayed with violin plots in blue, median counts of each sample are shown as white dots with black bars representing standard deviation. Data from 4 samples from Fos-TRAP^−^ cells and 3 samples from Fos-TRAP+ cells showed similar count distributions. Each sample contained cDNA from 150 neurons from each mouse. **(B)** Dispersion estimation of the normalized counts of samples at 3 days post-exercise was analyzed using the DEseq2 package. Local fit was used for the estimation, which showed that data dispersion estimated from raw counts (gene-est shown in black dots) fit well with the statistical model calculated from the DEseq2 package (red line). Normalized data (blue dots) were used for differential expression comparison.

**Fig. S5.**
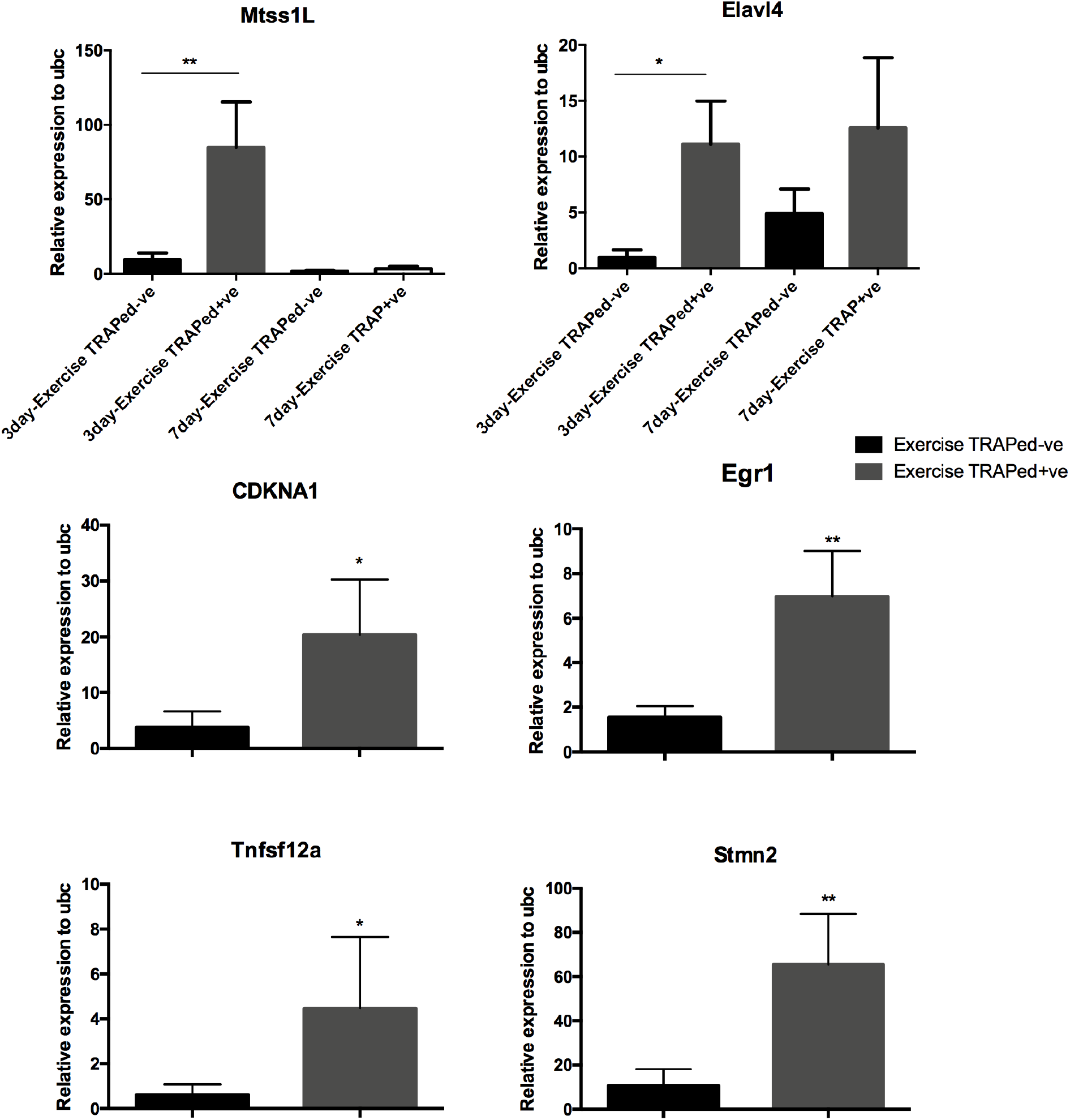
Real-time qPCR confirmation from the top 10 enriched genes in exercise-TRAPed cells at 3 and 7 days post-exercise. **(A)** Summary Real-Time qPCR confirmation of Mtss1L and Elavl4 exercise-induced transcripts in 3 and 7-day post-exercise samples using Smartseq2 amplified cDNA (exercise-TRAPed^−^ (control) n = 4 mice; exercise-TRAPed^+^ n = 3 mice). mRNA expression was normalized to neuronal internal control gene ubc. Statistical analysis was performed using one-way ANOVA with multiple comparisons and t-test *P<0.05. **(B)** mRNA expression of p21(CDKN1A), Egr1, Tnfsf12a, Stmn2 in the 3-day post-exercise samples (exercise TRAPed^−^ (control) n = 4 exercise TRAPed^+^ n = 3.). mRNA expression was normalized to neuronal internal control gene ubc. Statistical analysis was performed using t-test assuming equal variation *P<0.05.

**Fig. S6.**
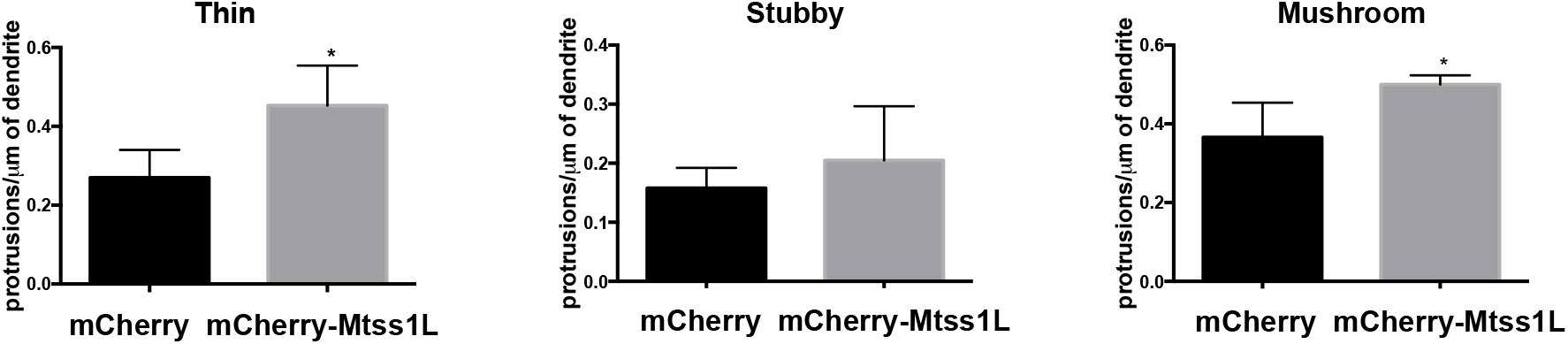
Effect of Mtss1L overexpression on dendritic spine subtypes of primary hippocampal neurons *in vitro.* Mtss1L overexpression significantly increased the density of both thin and mushroom dendritic spines, whereas there was no difference in the density of stubby spines (n=3, unpaired t-test, p<0.05).

**Fig. S7.**
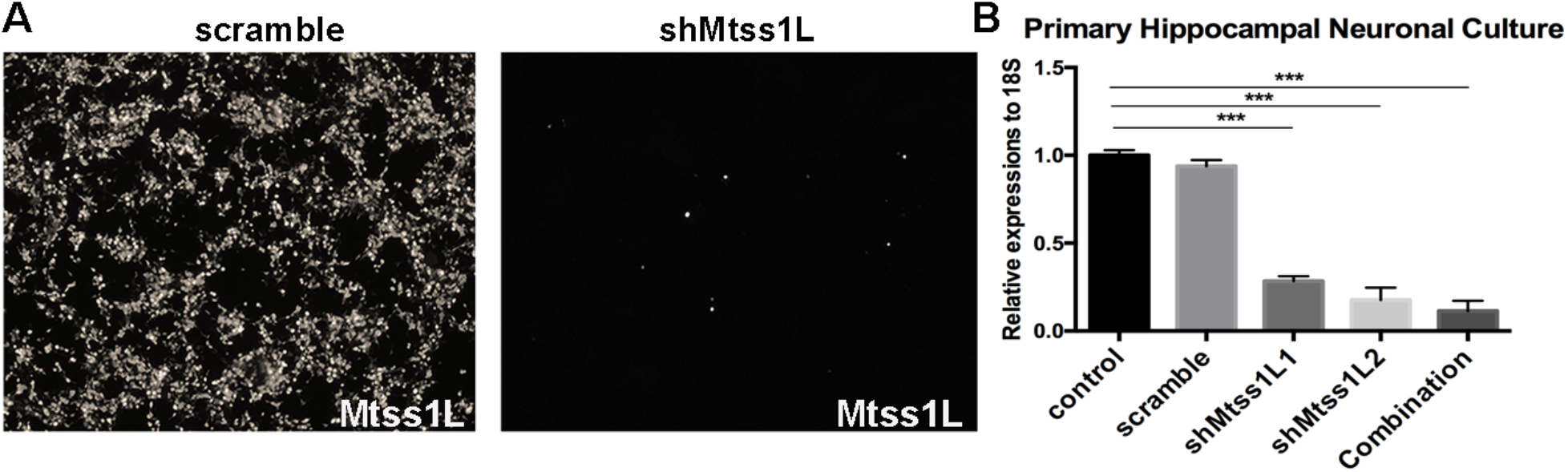
Mtss1L shRNA knockdown efficiency. **(A)** shRNA knockdown efficiency was first screened using HEK293T cells overexpressing Mtss1L after transduction with lentiviral construct FUGW-mcherry-T2A-Mtss1L. Knockdown efficiency of the overexpression construct was confirmed by fluorescence microscopy. **(B)** Primary hippocampal cultures were transduced with 10^^5^ particles of one or both short hairpin shRNAs against Mtss1L (designated L1 and L2) or shMtss1Lscramble lentivirus, and then treated for 7 d.i.v. with media supplemented by BDNF (25ng/ml). The shScramble virus was ineffective, whereas the combination of the two short hairpins was the most effective. Mtss1L shRNA knockdown efficiency was analyzed by RT-qPCR for Mtss1L expression, using 18S as the housekeeping/control gene (n=3 biological replicates, one way ANOVA with Dunnett’s multiple comparison, p<0.001).

**Fig. S8.**
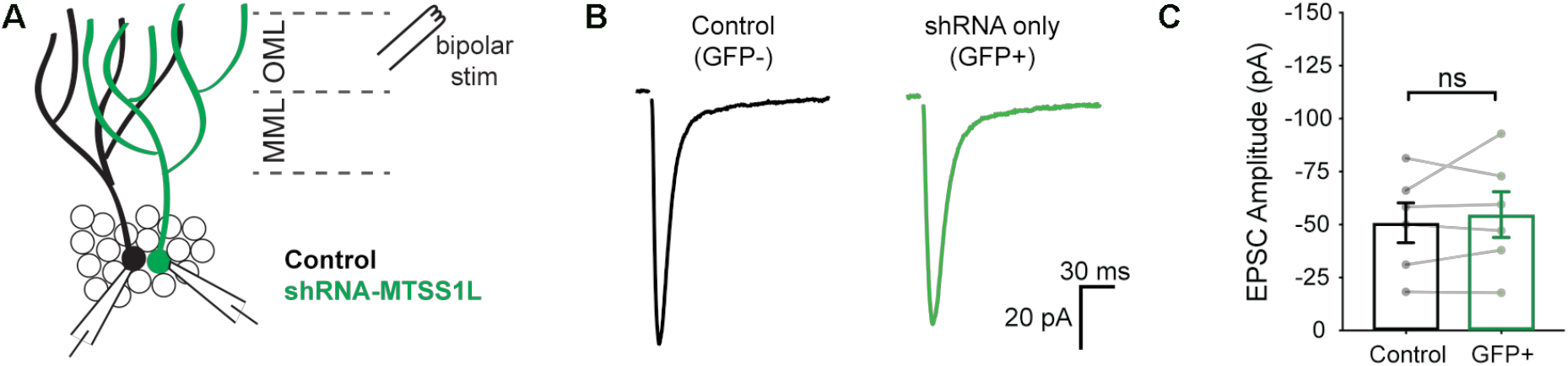
shRNA-Mtss1L did not alter evoked EPSCs in granule cells that were not exercise-TRAPed. **(A)** Schematic of paired dentate granule cell recordings. EPSCs were evoked with a bipolar stimulation electrode in the OML. **(B)** Representative EPSCs from a paired control granule cell (GFP^−^, black trace) and an adjacent Mtss1L shRNA only (GFP/tdT^−^) granule cell show similar amplitudes. Stimulation artifacts were blanked. **(C)** EPSC amplitudes in paired recordings were unaffected by shRNA-Mtss1L expression in granule cells that were not exercise-TRAPed, indicating that knockdown of Mtss1L had no effect unless the granule cell was exercise-activated (paired t-test, p = 0.4805).

**Table S1:**
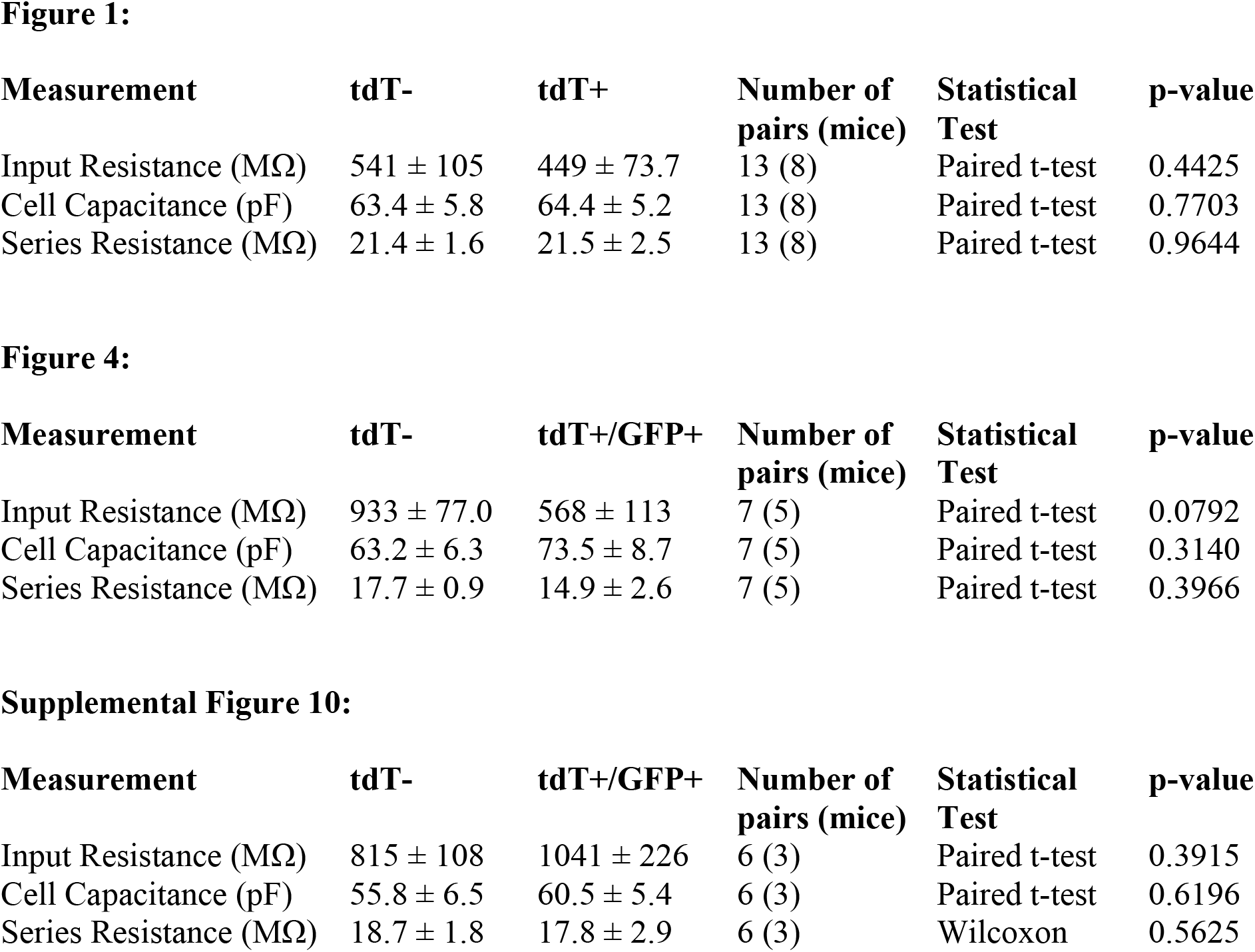
Intrinsic properties of granule cells during paired recordings.

| Measurement | tdT- | tdT+ | Number of pairs (mice) | Statistical Test | p-value |
| --- | --- | --- | --- | --- | --- |
| Input Resistance (MΩ) | 541 ± 105 | 449 ± 73.7 | 13 (8) | Paired t-test | 0.4425 |
| Cell Capacitance (pF) | 63.4 ± 5.8 | 64.4 ± 5.2 | 13 (8) | Paired t-test | 0.7703 |
| Series Resistance (MΩ) | 21.4 ± 1.6 | 21.5 ± 2.5 | 13 (8) | Paired t-test | 0.9644 |

**Figure 4:**
| Measurement | tdT- | tdT+/GFP+ | Number of pairs (mice) | Statistical Test | p-value |
| --- | --- | --- | --- | --- | --- |
| Input Resistance (MΩ) | 933 ± 77.0 | 568 ± 113 | 7 (5) | Paired t-test | 0.0792 |
| Cell Capacitance (pF) | 63.2 ± 6.3 | 73.5 ± 8.7 | 7 (5) | Paired t-test | 0.3140 |
| Series Resistance (MΩ) | 17.7 ± 0.9 | 14.9 ± 2.6 | 7 (5) | Paired t-test | 0.3966 |

| Measurement | tdT- | tdT+/GFP+ | Number of pairs (mice) | Statistical Test | p-value |
| --- | --- | --- | --- | --- | --- |
| Input Resistance (MΩ) | 815 ± 108 | 1041 ± 226 | 6 (3) | Paired t-test | 0.3915 |
| Cell Capacitance (pF) | 55.8 ± 6.5 | 60.5 ± 5.4 | 6 (3) | Paired t-test | 0.6196 |
| Series Resistance (MΩ) | 18.7 ± 1.8 | 17.8 ± 2.9 | 6 (3) | Wilcoxon | 0.5625 |

Table S2 & S3 are attached as Auxiliary Files.

**Table S4:**
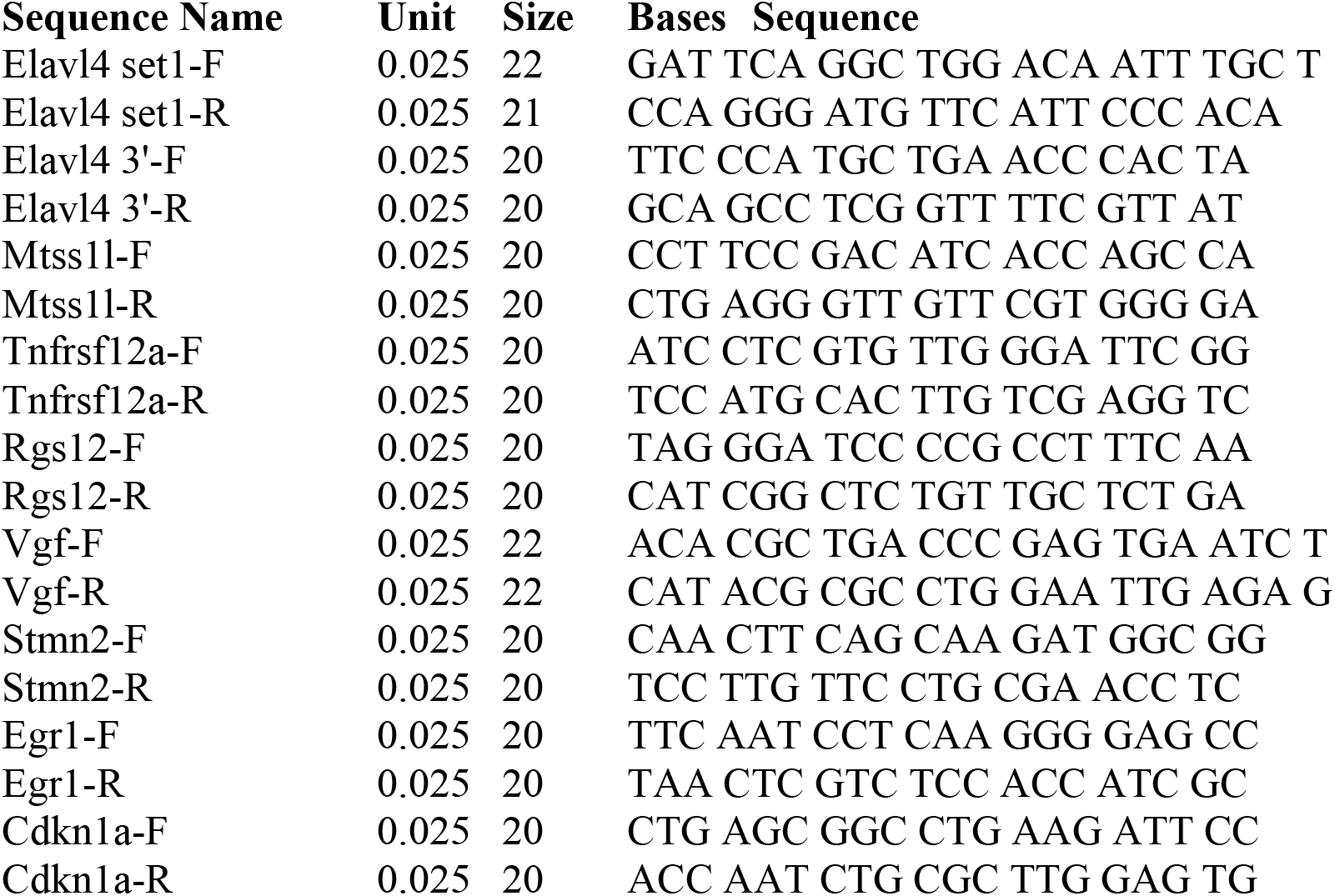
List of primers used for RT-qPCR analysis.

## References

1. L. Mandolesi et al., Effects of Physical Exercise on Cognitive Functioning and Wellbeing: Biological and Psychological Benefits. Front Psychol 9, 509 (2018).

2. D. Saraulli, M. Costanzi, V. Mastrorilli, S. Farioli-Vecchioli, The Long Run: Neuroprotective Effects of Physical Exercise on Adult Neurogenesis from Youth to Old Age. Curr Neuropharmacol 15, 519–533 (2017).

3. C. Vivar, M. C. Potter, H. van Praag, All about running: synaptic plasticity, growth factors and adult hippocampal neurogenesis. Curr Top Behav Neurosci 15, 189–210 (2013).

4. M. W. Voss, C. Vivar, A. F. Kramer, H. van Praag, Bridging animal and human models of exercise-induced brain plasticity. Trends Cogn Sci 17, 525–544 (2013).

5. M. W. Jung, B. L. McNaughton, Spatial selectivity of unit activity in the hippocampal granular layer. Hippocampus 3, 165–182 (1993).

6. J. K. Leutgeb, S. Leutgeb, M. B. Moser, E. I. Moser, Pattern separation in the dentate gyrus and CA3 of the hippocampus. Science 315, 961–966 (2007).

7. W. Severa, O. Parekh, C. D. James, J. B. Aimone, A Combinatorial Model for Dentate Gyrus Sparse Coding. Neural Comput 29, 94–117 (2017).

8. L. Bolz, S. Heigele, J. Bischofberger, Running Improves Pattern Separation during Novel Object Recognition. Brain Plast 1, 129–141 (2015).

9. D. J. Creer, C. Romberg, L. M. Saksida, H. van Praag, T. J. Bussey, Running enhances spatial pattern separation in mice. Proc Natl Acad Sci U S A 107, 2367–2372 (2010).

10. M. Uda, M. Ishido, K. Kami, M. Masuhara, Effects of chronic treadmill running on neurogenesis in the dentate gyrus of the hippocampus of adult rat. Brain Res 1104, 64–72 (2006).

11. R. Perini, M. Bortoletto, M. Capogrosso, A. Fertonani, C. Miniussi, Acute effects of aerobic exercise promote learning. Sci Rep 6, 25440 (2016).

12. J. C. Basso, W. A. Suzuki, The Effects of Acute Exercise on Mood, Cognition, Neurophysiology, and Neurochemical Pathways: A Review. Brain Plast 2, 127–152 (2017).

13. E. L. Hargreaves, G. Rao, I. Lee, J. J. Knierim, Major dissociation between medial and lateral entorhinal input to dorsal hippocampus. Science 308, 1792–1794 (2005).

14. D. Yoganarasimha, G. Rao, J. J. Knierim, Lateral entorhinal neurons are not spatially selective in cue-rich environments. Hippocampus 21, 1363–1374 (2011).

15. T. Hafting, M. Fyhn, S. Molden, M. B. Moser, E. I. Moser, Microstructure of a spatial map in the entorhinal cortex. Nature 436, 801–806 (2005).

16. F. Sargolini et al., Conjunctive representation of position, direction, and velocity in entorhinal cortex. Science 312, 758–762 (2006).

17. A. E. West, M. E. Greenberg, Neuronal activity-regulated gene transcription in synapse development and cognitive function. Cold Spring Harb Perspect Biol 3, (2011).

18. W. Akamatsu et al., The RNA-binding protein HuD regulates neuronal cell identity and maturation. Proc Natl Acad Sci U S A 102, 4625–4630 (2005).

19. M. W. Jones et al., A requirement for the immediate early gene Zif268 in the expression of late LTP and long-term memories. Nat Neurosci 4, 289–296 (2001).

20. J. Saarikangas et al., ABBA regulates plasma-membrane and actin dynamics to promote radial glia extension. J Cell Sci 121, 1444–1454 (2008).

21. W. C. Skarnes et al., A conditional knockout resource for the genome-wide study of mouse gene function. Nature 474, 337–342 (2011).

22. J. C. Dawson, J. A. Legg, L. M. Machesky, Bar domain proteins: a role in tubulation, scission and actin assembly in clathrin-mediated endocytosis. Trends Cell Biol 16, 493–498 (2006).

23. S. Vaynman, Z. Ying, F. Gomez-Pinilla, Hippocampal BDNF mediates the efficacy of exercise on synaptic plasticity and cognition. Eur J Neurosci 20, 2580–2590 (2004).

24. C. D. Wrann et al., Exercise induces hippocampal BDNF through a PGC-1alpha/FNDC5 pathway. Cell Metab 18, 649–659 (2013).

25. G. G. Gross et al., Recombinant probes for visualizing endogenous synaptic proteins in living neurons. Neuron 78, 971–985 (2013).

26. H. van Praag, B. Christie, Tracking Effects of Exercise on Neuronal Plasticity. Brain Plast 1, 3–4 (2015).

27. H. van Praag, Neurogenesis and exercise: past and future directions. Neuromolecular Med 10, 128–140 (2008).

28. L. S. Overstreet et al., A transgenic marker for newly born granule cells in dentate gyrus. J Neurosci 24, 3251–3259 (2004).

29. J. Saarikangas et al., MIM-Induced Membrane Bending Promotes Dendritic Spine Initiation. Dev Cell 33, 644–659 (2015).

30. P.K. Mattila et al., Missing-in-metastasis and IRSp53 deform PIP2-rich membranes by an inverse BAR domain-like mechanism. J. Cell Biol. 7, 953–964 (2007).

31. C. Charrier et al., Inhibition of SRGAP2 function by its human-specific paralogs induces neoteny during spine maturation. Cell 149, 923–935 (2012).

32. J. Choi et al., Regulation of dendritic spine morphogenesis by insulin receptor substrate 53, a downstream effector of Rac1 and Cdc42 small GTPases. J Neurosci 25, 869–879 (2005).

33. A. Tsao et al., Integrating time from experience in the lateral entorhinal cortex. Nature 561, 57–62 (2018).

34. E. V. van Dongen, I. H. P. Kersten, I. C. Wagner, R. G. M. Morris, G. Fernandez, Physical Exercise Performed Four Hours after Learning Improves Memory Retention and Increases Hippocampal Pattern Similarity during Retrieval. Curr Biol 26, 1722–1727 (2016).

35. M. L. Seibenhener, M. W. Wooten, Isolation and culture of hippocampal neurons fromprenatal mice. J Vis Exp, (2012).

36. H. Ito, R. Morishita, I. Iwamoto, K. Nagata, Establishment of an in vivo electroporation method into postnatal newborn neurons in the dentate gyrus. Hippocampus 24, 1449–1457 (2014).

37. W. D. Hendricks, Y. Chen, A. L. Bensen, G. L. Westbrook, E. Schnell, ShortTerm Depression of Sprouted Mossy Fiber Synapses from Adult-Born Granule Cells. J Neurosc 37, 5722–5735 (2017).

38. N. I. Woods et al., Preferential Targeting of Lateral Entorhinal Inputs onto Newly Integrated Granule Cells. JNeurosci 38, 5843–5853 (2018).

